# Effects of age and hearing loss on perceptual and physiological measures of temporal envelope processing and spatial release from speech-on-speech masking

**DOI:** 10.1101/2020.09.03.281717

**Authors:** Chhayakanta Patro, Heather A. Kreft, Magdalena Wojtczak

## Abstract

Older adults often experience difficulties understanding speech in adverse listening conditions. These difficulties are partially attributed to auditory temporal-processing deficits associated with aging even in the absence of hearing loss. The aim of this study was to assess effects of age and hearing loss on temporal envelope processing and speech-on-speech masking. Listeners with normal and near-normal hearing across a wide age range (20 to 66 years) were tested using a series of psychophysical (amplitude-modulation detection, gap detection, and interaural-envelope-phase discrimination), physiological (electroencephalographic envelope-following responses), speech perception (spatial release from masking), and cognitive (processing speed) measures. Results showed that: (i) psychophysical measures of monaural and binaural envelope processing and neural measures of envelope processing are not affected by aging after accounting for audiometric hearing loss, (ii) behavioral gap-detection thresholds decline with age, (iii) aging results in a reduction of spatial release from masking, even as speech intensity is amplified in the region of hearing loss, (iv) aging is associated with poorer measures of cognitive function. Although age significantly contributed to a decline in spatial release from speech-on-speech masking, individual differences in envelope processing and in scores from nonauditory cognitive tests used in this study were not significant predictors of speech performance.

**Highlights:**

- Age per se does not affect psychophysical and physiological measures of monaural amplitude-modulation processing.
- Age does not affect the ability to detect interaural disparities in envelope timing between the ears.
- Gap detection thresholds degrades with age even after hearing thresholds are statistically accounted for.
- Age, independent of hearing thresholds, can substantially reduce spatial release from masking.
- Cognitive ability declines with age. However, such declines do not necessarily cause deficits in spatial release from masking.

## 1 Introduction

Speech perception difficulties in noisy acoustic environments are very common among aging individuals (CHABA, 1988). These difficulties occur even in the absence of hearing loss (Dubno et al., 2002; Frisina et al., 1997; Pichora-Fuller et al., 2003), indicating that factors unrelated to cochlear damage are also important contributors. One of the main factors shown to affect speech understanding is the fidelity of temporal processing of suprathreshold sounds. There are two main types of temporal information carried by acoustic stimuli, the temporal fine structure and the envelope. The temporal fine-structure processing requires phase locking of neural responses to fast changes in sound pressure in the stimulus and it has been shown to be affected by age and cochlear hearing loss (Füllgrabe et al., 2015; Hopkins et al., 2011; Hopkins et al., 2008; Lorenzi et al., 2006; Moore, 2016; Strelcyk et al., 2009). The temporal envelope is defined by relatively slow changes in stimulus amplitude over time and it appears to be generally unaffected by hearing loss (Bacon et al., 1985; Moore et al., 2001). However, effects of age on temporal-envelope processing are less clear. While some studies have reported no differences in the processing of amplitude modulation (AM) between older and younger normal-hearing adults (Boettcher et al., 2001; Grose et al., 2019; Schoof et al., 2014), others found that envelope processing declines with age, particularly at fast modulation rates (Kumar et al., 2011; Purcell et al., 2004). Age-related deficits in temporal-envelope processing have been consistently observed in animal models (Herrmann et al., 2017; Parthasarathy et al., 2018a; Parthasarathy et al., 2018b), suggesting that similar declines could occur in aging humans.

Auditory information contained in the temporal envelope has been shown to play an important role for speech understanding (Goossens et al., 2018; Rosen, 1992; van Tasell et al., 1987). Physiological measures have shown that the fidelity of speech-envelope representation in neural responses is strongly related to speech perception (Ahissar et al., 2001; Goossens et al., 2018; Peelle et al., 2012). The important role of robust temporal-envelope representation for understanding speech in noise is also supported by studies showing that speech intelligibility can be successfully predicted based solely on the target-to-masker envelope power ratio (Jørgensen et al., 2011; Jørgensen et al., 2013; Jørgensen et al., 2015).

Until about a decade ago, age-related deficits in auditory temporal processing and in perception of speech in noise have been often attributed to age-related changes at central sites of the auditory pathways and to general decline in cognitive function (Akeroyd, 2008; Dubno et al., 2002; Frisina et al., 1997; Pichora-Fuller et al., 2003; Pichora-Fuller et al., 1995; Rajan et al., 2008; Saunders et al., 1992). However, a discovery of diffuse loss of auditory-nerve synapses in mice due to noise exposure (Kujawa et al., 2009) and healthy aging (Sergeyenko et al., 2013) provided a base for a new hypothesis that some decline in speech recognition in noisy backgrounds could originate from dysfunction at the peripheral level of the auditory system. According to this hypothesis, older individuals with normal or near-normal hearing and no signs of cognitive decline may experience difficulties understanding speech in noise because of a reduced fidelity of temporal representation of speech in auditory-nerve responses. The findings from animal data have motivated a large number of studies searching for evidence of temporal-processing deficits in humans that could be attributed to cochlear synaptopathy. The studies, which are mostly based on measures obtained from young normal-hearing adults with different histories of noise exposure, have provided conflicting results. While some of the studies report findings that support the hypothesis based on cochlear synaptopathy (Grose et al., 2017; Liberman et al., 2016; Stamper et al., 2015; Valderrama et al., 2018), others show no evidence for the presence of temporal-processing deficits expected from cochlear synaptopathy (Grinn et al., 2017; Guest et al., 2017b; Prendergast et al., 2017a; Prendergast et al., 2017b; Spankovich et al., 2014; Yeend et al., 2017).

A smaller number of studies have included older normal or near-normal hearing individuals when exploring different proxy measures of cochlear synaptopathy (Mepani et al., 2020; Prendergast et al., 2019). The main reason for limiting the study samples to young adults is that older individuals often have higher audiometric thresholds even when the thresholds are within the limits of “normal hearing” as defined by ANSI standards (ANSI, 1996). Higher hearing thresholds make it hard to tease apart effects of age and hearing sensitivity. Direct evidence from human temporal bones shows that loss of auditory-nerve axons becomes increasingly severe with advancing age (Wu et al., 2019), even as some of the observed loss might be due to combined effects of age and noise exposure. Prendergast et al. (2019) added a population of older adults to a large group of young normal-hearing individuals tested in their earlier studies (Prendergast et al., 2017a; Prendergast et al., 2017b). Combining younger and older adults resulted in a large total sample of individuals with ages in a range of 18-60 years. Prendergast et al. (2019) used a few measures of temporal processing, including amplitude-modulation (AM) detection thresholds, interaural envelope-phase-discrimination (envIPD) thresholds, and the magnitude of electroencephalographic (EEG) envelope following responses (EFRs). None of the measures was significantly correlated with age. However, significant age effects on behavioral measures of AM processing and on EFRs have been found in other studies (e.g., Dimitrijevic et al., 2016; Leigh-Paffenroth et al., 2006; Purcell et al., 2004). These age-related deficits were observed for high (≥ ~100 Hz) but not for lower modulation rates (Grose et al., 2009; Purcell et al., 2004). Measures of temporal envelope processing are often significantly correlated with hearing sensitivity (e.g., Grose et al., 2009; Prendergast et al., 2019; Purcell et al., 2004) suggesting that some of the reported age effects may have been related to deficits in cochlear function rather than aging per se.

In this study, a battery of psychophysical, electrophysiological, and speech perception measures was used to assess effects of age on temporal envelope processing and on spatial release from masking (SRM) for speech presented in two-talker babble. Our targeted population were individuals with ages spanning a wide range (18-70 yrs) and with normal hearing. However, we found a relatively small number of older listeners with hearing thresholds matching those of our youngest participants. Because of that, individuals with mild-to-moderate high-frequency hearing loss were included in the study and the effects of hearing sensitivity were statistically controlled for in data analyses. Based on the temporal bone study by Wu et al. (2019), we hypothesized that deficits in temporal processing will be observed with increasing age even after the effects of hearing loss have been accounted for. If present, such deficits would be consistent with the hypothesis that cochlear synaptopathy, which likely coexists with cochlear hearing loss, adds to difficulties with speech-in-noise perception and auditory scene analysis in the aging population.

## 2 Psychophysical measures of temporal-envelope processing

### 2.1 Participants

A total of 61 adults (22 males, 38 females, 1 other) with the mean age of 45.6 years (range 20 - 66 yrs) were recruited from the University of Minnesota and surrounding community. For the purpose of investigating age effects, all data analyses in this study were performed with the participants divided into three age groups, young (20-35 yrs), middle (36-50 yrs), and older (51-66 yrs) and all collected measures were compared across these age groups. The reason for comparing mean data across the three age groups was to obtain a rough estimate of an age bracket during which age-related changes in auditory processing begin to significantly affect the specific measures used in this study. For each participant, audiometric thresholds were measured at octave frequencies between 250 and 8000 Hz, using a calibrated audiometer (Madsen Conera, GN Otometrics) to evaluate their hearing status. Normal hearing was defined by pure-tone air-conduction thresholds less than or equal to 20 dB HL. All participants had normal thresholds at frequencies ≤ 2000 Hz in both ears, but some older individuals had mild hearing loss at the two highest frequencies, 4000 and 8000 Hz. All participants had approximately symmetric hearing (across-ear differences ≤ 10 dB) at all audiometric frequencies. In addition to the standard clinical audiometry, hearing thresholds were also measured using a two-alternative forced-choice (2AFC) procedure with a two-down, one-up adaptive-tracking rule that estimates the 70.7% correct point on the psychometric function (Levitt, 1971). The measurements obtained with the adaptive tracking were performed using the AFC software package (Ewert, 2013) using MATLAB (The Mathworks, Natick, MA). The 2AFC procedure was used to obtain a fine-grained unbiased measure of hearing thresholds for better statistical control of effects of hearing sensitivity. The steps in the adaptive tracking were 8 dB for the first two reversals, 4 dB for the subsequent two reversals, and 2 dB for the final eight reversals. The tone level and the final eight reversals were averaged to calculate threshold. Two estimates of hearing threshold, in dB sound pressure level (dB SPL), were averaged to obtain the final threshold estimate at each audiometric frequency. Thresholds averaged across the three age groups are shown by different symbols (and lines connecting them) in Fig. 1.

**Fig. 1.**
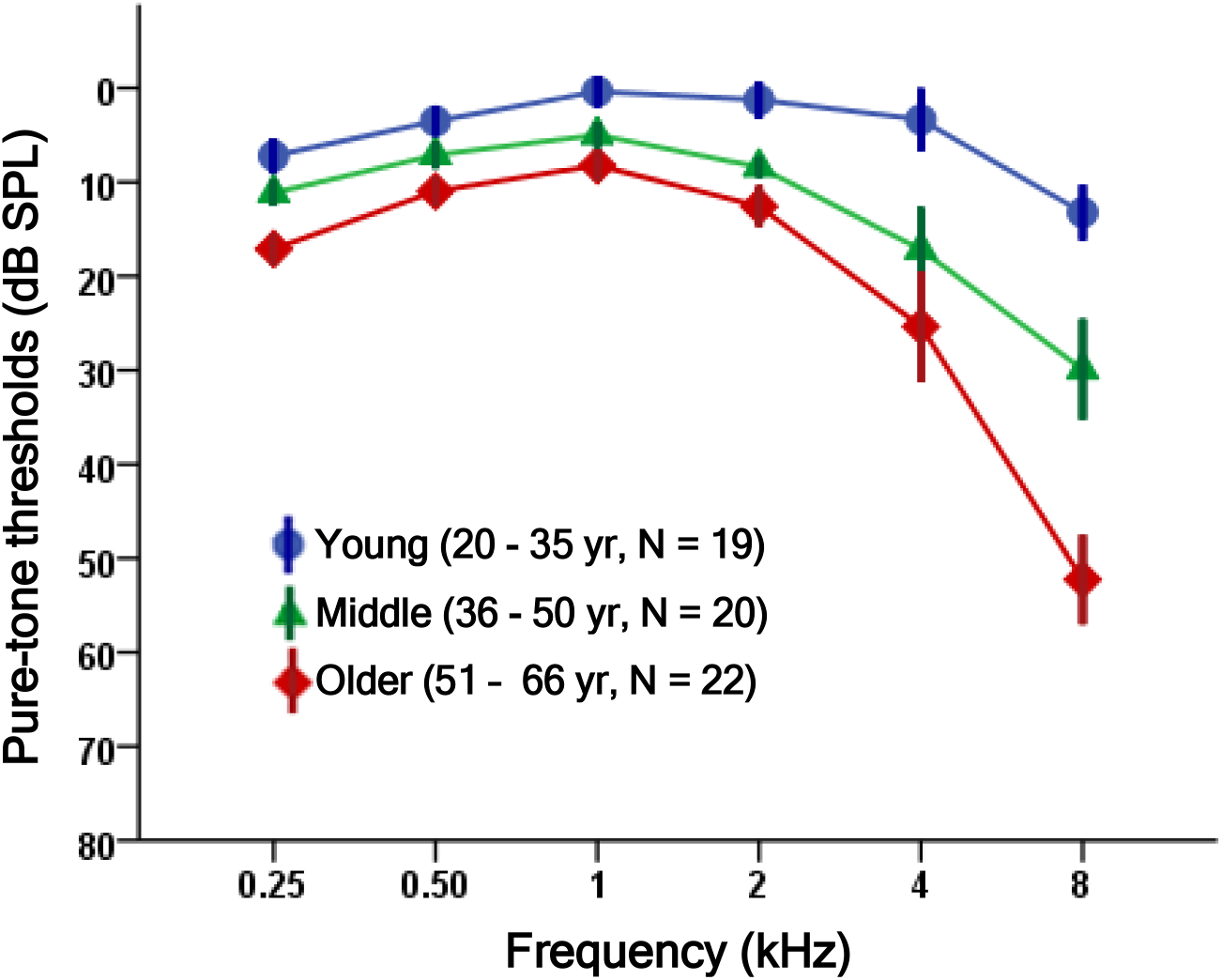
Hearing thresholds, measured in dB SPL, using a 2AFC adaptive-tracking method for three age groups specified in the legend. The error bars indicate ± one standard error (SE) of the mean.

Despite similar audiometric thresholds across the three groups at frequencies ≤ 2 kHz, thresholds measured with the 2AFC adaptive-tracking procedure averaged within each age group show progressive loss of hearing sensitivity with increasing age at all frequencies between 0.25 and 8 kHz, with the sensitivity curves increasingly diverging at frequencies above 2 kHz. A mixed-design analysis of variance (ANOVA), with the tone frequency as the within-subjects factor and age group as the between-subjects factor, showed a significant effect of frequency [F(5, 290) = 43.2, p < 0.001], a significant effect of age group [F(2, 58) = 17.7, p < 0.001], and a significant interaction between frequency and age group [F(10, 290) = 5.7, p <0.001]. Post hoc pairwise comparisons showed significant differences in hearing sensitivity between young and middle age groups (p = 0.012), young and older groups (p < 0.001), and middle and older groups (p = 0.011).

In addition to hearing sensitivity, tympanometry was performed in each listener to ensure normal middle-ear function (type-A tympanogram) using a calibrated tympanometer (MT10, Interacoustics, DK). Not all individuals, who consented to participate in this study, completed the entire protocol. Table 1 describes the sample size, mean age of the sample, the average of hearing thresholds in dB SPL used as covariates in the statistical analyses for each experiment (described below), and a range of these average hearing thresholds across participants in a sample for each task. The average hearing thresholds represent mean thresholds across the range from 2 to 8 kHz, with the exception of the speech experiment, for which the average was calculated over the whole audiometric range (0.25 – 8 kHz). A total of 27 recruited individuals were able to participate in all the experiments. Participants received monetary compensation after each experimental session. Each participant provided written informed consent before testing began and the experimental protocols were approved by the Institutional Review Board at the University of Minnesota.

**Table 1.** The sample size, the mean age, the mean hearing threshold (average across 2 – 8 kHz for all tasks except speech, for which the average was calculated across 0.25 – 8 kHz), the range of average hearing thresholds, for listeners participating in each task in this study (listed in column 1). The age range of participants (20 – 66 yrs) was the same for each task.

| **Experimental task** | **Sample size** | **Mean age (years)** | **Mean AVG_HT (dB SPL)** | **AVG_HT range (dB SPL)** |
| --- | --- | --- | --- | --- |
| **AM Detection** | 61 (38 females) | 45.64 | 18.1 | -18 – 68.9 |
| **Gap Detection** | 53 (32 females) | 42.84 | 20.1 | -17.7 – 73.3 |
| **envITD** | 27 (20 females) | 43.61 | 19.8 | 1 – 66.5 |
| **Speech** | 58 (36 females) | 45.43 | 19.8 | -17.3 – 55.5 |
| **EFR** | 45 (33 females) | 44.1 | 18.0 | -18 – 51.5 |
| **All tasks** | 27 (20 females) | 53.61 | 19.8 | 1 – 66.5 |

### 2.2 Stimuli and procedures

#### 2.2.1 Detection of AM for a noise carrier

Thresholds for detecting sinusoidal AM in a Gaussian-noise carrier were measured for three modulation rates, 100, 300, and 600 Hz. These modulation rates were chosen because previous studies showed age effects on AM responses for fast, but not for slower rates (Grose et al., 2009; Purcell et al., 2004). The carrier had a bandwidth from 0.1 to 10 kHz. A 2AFC adaptive-tracking procedure combined with a three-down, one-up tracking rule was used to obtain the threshold estimate corresponding to the 79.4 % correct point on the psychometric function (Levitt, 1971). In one interval, selected at random, the noise was modulated throughout its duration while the other interval contained an unmodulated noise. The stimuli in the two intervals were scaled to have equal root-mean-squared amplitudes to eliminate loudness cues (Viemeister, 1979) and were presented at 70 dB SPL. The noise carrier had a duration of 250 ms including 10-ms onset/offset ramps, and the two observation intervals were separated by 300 ms of silence. The observation intervals were marked by flashing colored boxes on a computer screen. In each trial, listeners were asked to choose the interval containing the AM and provide their response by a keypress or via a mouse click. Visual feedback indicating the correct response was provided after each trial. The tracking variable was the modulation depth expressed in dB [20 log (*m*)], where *m* is the modulation index with a value between 0 and 1. The modulation depth was changed in 4-dB steps for the first two reversals, 2-dB steps for the subsequent two reversals, and 1-dB steps for the final eight reversals. AM detection threshold from a single run was calculated by averaging modulation depths (in dB) at the last eight reversals. Thresholds from three runs were averaged to compute the final threshold estimate. When the adaptive-tracking procedure called for a modulation depth greater than 0 dB (100% AM), the track was aborted and another measure was started. This happened for 10 listeners in one of the three conditions (usually for the 600-Hz modulation rate). All 10 listeners were able to successfully complete the track after one aborted run.

#### 2.2.2 Gap detection for tones in noise

Gap detection thresholds were measured at two frequencies, 2 and 4 kHz, for tones embedded in one-octave Gaussian noise. For each tone, the noise was geometrically centered at the tone’s frequency. The two frequencies were selected because at the lower frequency (2 kHz) no listeners had clinical hearing loss, and at the other frequency (4 kHz) some older participants exhibited a mild loss of hearing sensitivity (i.e., audiometric thresholds ≥ 20 dB HL). The levels of the tones and noise bands were 75 and 65 dB SPL, respectively, resulting in a signal-to-noise ratio (SNR) of 10 dB. A three-interval, three-alternative forced-choice (3AFC) procedure combined with a three-down, one-up adaptive tracking technique was used to measure thresholds corresponding to the 79.4 % correct point on the psychometric function (Levitt, 1971). The tonal markers before and after the gap were 250-ms long including 1-ms hanning onset and offset ramps. The noise was 700-ms long including 10-ms ramps. In each observation interval, the first marker started 50 ms after the noise onset. In the non-signal intervals, the markers were presented in sequence with a 0-ms gap between the offset of the first marker and the onset of the second marker. In the signal interval, a silent gap was inserted between the markers. The tracking variable was the gap duration expressed in logarithmic units of 10log(Δ*t*), where *Δ*t** was the gap duration in milliseconds. The gap duration was varied in steps of 3 logarithmic units for the first two reversals, 1.5 units for the subsequent two reversals, and 0.75 units for the final eight reversals. A limit was set for the gap duration to abort the track when the adaptive procedure called for a duration that would cause the post-gap marker to extend beyond the duration of the noise masker. None of the listeners reached that limit. Threshold from a single run was calculated by averaging the logarithmically transformed gap durations at the last eight reversals. The final estimate was obtained by averaging thresholds from three runs.

#### 2.2.3 Interaural envelope time difference (envITD)

Sensitivity to envITDs was measured using a 2AFC procedure combined with a two-down, one-up adaptive tracking technique estimating threshold at the 70.7% correct point on the psychometric function (Levitt, 1971). The measurements were performed at two carrier frequencies, 2 and 4 kHz, and for two rates, 40 and 100 Hz, of full (100%) sinusoidal AM at each carrier frequency. In each interval, a sequence of four 500-ms AM tonal carriers separated by 20-ms silent gap was presented to both ears. The carrier duration included 100-ms hanning onset and offset ramps. In the signal interval, selected at random, the first and third tones had AM with the same phase in the two ears (diotic AM tones), while the AM in the second and fourth tones had an AM phase difference between the two ears, resulting in envITD (an ABAB sequence similar to that used by Hopkins et al., 2010). Because relatively high modulation rates were used, for suprathreshold envITDs the stimulus was perceived as moving between narrowly centered intracranial sound image to a broader image spreading toward both sides of the head over the course of four tone bursts in the observation interval. The non-signal interval consisted of four diotically presented AM tones (AAAA sequence), resulting in a fixed (centered) intracranial sound image. The envITD thresholds were measured in quiet and in the presence of a notched-noise masker. The noise had a bandwidth from 0.1 to 10 kHz and contained a spectral notch with a width of 0.2 *x* carrier frequency, geometrically centered on the carrier frequency. The notched noise started 100 ms before the first observation interval and ended 100 ms after the end of the second observation interval. Different samples of noise were generated for each trial separately for the left and right ear. The two observation intervals were separated by a 500-ms gap. For thresholds measured in quiet the gap contained silence, and for thresholds in noise, the notched noise continued throughout the gap between the observation intervals. The carrier level was 70 dB SPL and the notched noise had an overall level of 50 dB SPL. Listeners were asked to choose the interval in which they perceived movement of the sound image in their head across the four stimulus presentations. The observation intervals were marked by flashing colored boxes and visual feedback indicating the correct response was provided after each trial. The adaptive tracking variable was a log-transformed interaural envelope phase difference. The track started with the phase difference set to log(π). The phase difference was changed in steps of log(1.25^2^) for the first four reversals and log(1.25) for the remaining eight reversals. Thresholds from a single run were estimated by averaging the log-transformed interaural envelope phase differences at the final eight reversals. Three single-run threshold estimates were averaged to calculate the final threshold. The log-transformed interaural phase difference at threshold was then converted to the interaural time difference, envITD.

For all the psychophysical tasks, listeners were seated in a single-walled sound-attenuating booth. The stimuli were generated in Matlab (Mathworks, Natick, MA) on a PC and were played using an E22 soundcard (LynxStudio, Costa Mesa, CA) with 24-bit resolution and a sampling rate of 48 kHz. The different experimental tasks were implemented using the AFC software package (Ewert, 2013) utilizing Matlab. For the monaural tasks, the stimuli were delivered to the left ear, and for the envITD task to both ears, through Sennheiser HD650 headphones (Sennheiser, Old Lyme, CT).

### 2.3 Results and discussion

#### 2.3.1 AM detection

A total sample of 61 listeners completed the AM-detection experiment. The mean thresholds for the three age groups are shown by different color bars in Fig. 2. As expected, the three groups exhibited an increase (worsening) of AM detection threshold with increasing modulation rate between 100 and 600 Hz, consistent with the low-pass characteristic of the temporal modulation transfer function for broadband noise carriers (Viemeister, 1979).

**Fig. 2.**
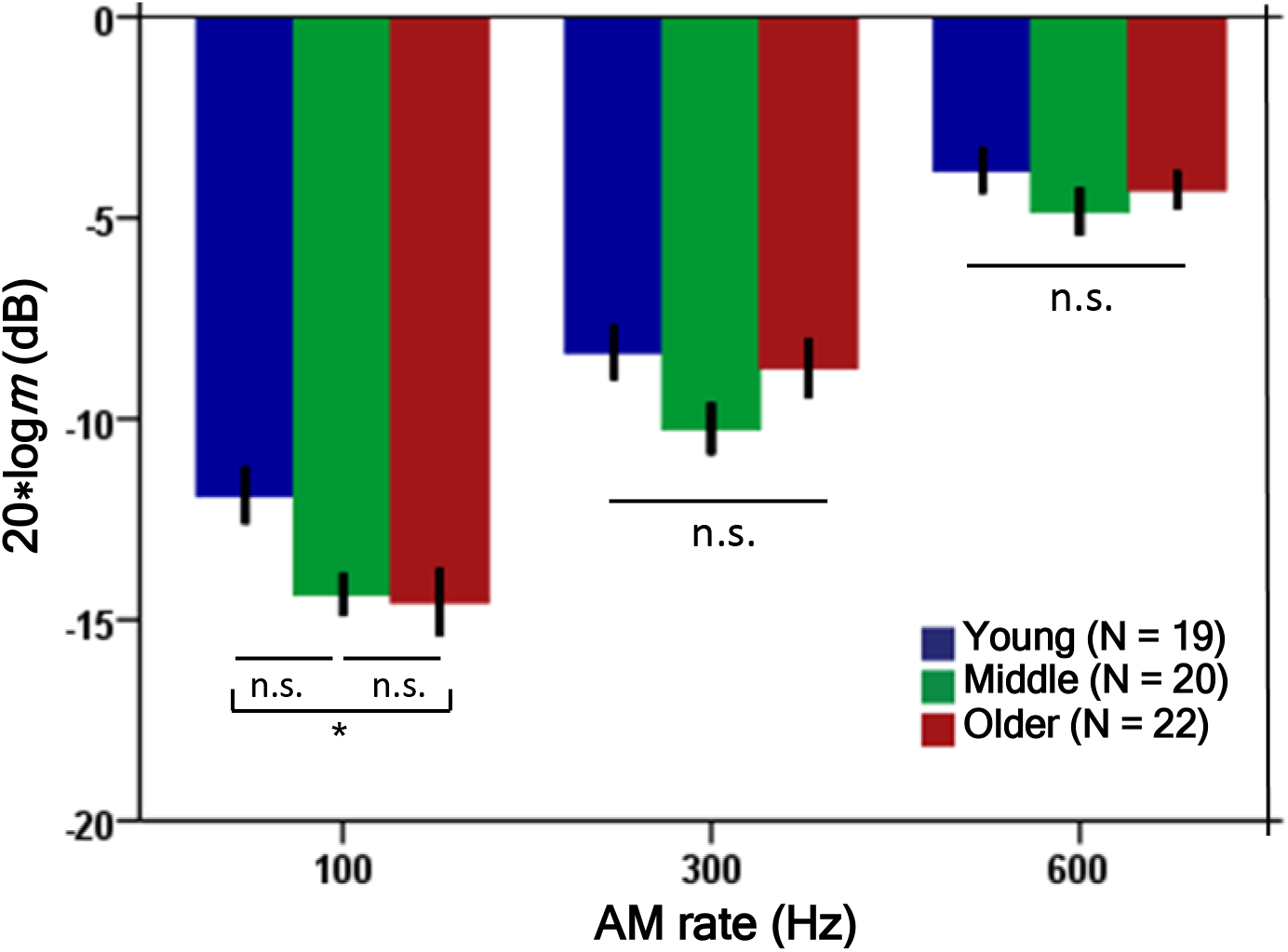
AM-detection thresholds for broadband noise carrier as a function of modulation rate, for three age groups. The error bars indicate ± 1 SE. The asterisk denotes a significant difference (* p < 0.05).

A mixed-design ANOVA was performed on AM-detection thresholds with the modulation rate as the within-subjects factor and the age group as the between-subjects factor. The average of hearing thresholds, measured in dB SPL, for frequencies of 2, 4, and 8 kHz (HFAVG) was used as a covariate. These frequencies were included in the covariate because hearing thresholds diverged between the three age groups in the high-frequency region and because high modulation rates are processed in auditory filters tuned to high frequencies. Because Mauchly’s test showed that sphericity assumption was violated, Greenhouse-Geisser correction was applied in reporting the results of the within-subjects effects. The ANOVA showed a significant effect of the modulation rate [F(1.6, 93.4) = 132.81, p < 0.001], a significant effect of HFAVG [F(1, 57) = 5.59, p = 0.022], but no significant effect of age group once the HFAVG had been controlled for [F(2, 57) = 1.55, p = 0.22]. No significant interactions were observed between the AM rate and HFAVG [F(1.6, 93.4) = 1.70, p = 0.187] or between the AM rate and age group [F(3.3, 93.4) = 2.10, p = 0.1]. Post hoc Game-Howell pairwise comparisons between the three age groups for each AM rate showed that AM-detection thresholds were significantly higher for the young than the middle age group (p = 0.026) for the AM rate of 100 Hz only. There were no other significant differences between age groups at this and the other two AM rates (p > 0.05).

The general lack of the effect of age group on the sensitivity to high-rate AM contrasts with significant age effects reported by Kumar et al. (2011). Their listeners’ age spanned a wider age range (20 – 85 yrs), and they reported a deterioration of AM-detection threshold with age between the youngest group (20-30 yrs) and the middle-aged group (41-50 yrs), with no further decline in performance for the older listeners (50 – 86 yrs) at a modulation rate of 200 Hz (the highest used in that study). It is unclear what accounts for the different finding in this study. AM-detection thresholds for the 100- and 300-Hz modulation rates (shown in Fig. 2) were lower for all age gups than those reported by Kumar et al. (2011) for the 200-Hz rate. In particular, the middle and older groups of participants in this study performed much better in the task than the listeners older than 41 yrs in Kumar et al. (2011), despite somewhat greater loss of hearing sensitivity and a lower level of the stimuli used here.

The lower (better) 100-Hz AM-detection threshold observed in the middle and older than in the young group of listeners (Fig.2, leftmost set of bars) could be due to outer hair cell (OHC) damage at basal sites of the cochlea of our older participants. This explanation is supported by the significant between-subjects effect of HFAVG and a lack of thereof for age group, as reported above. OHC loss is associated with broadening of cochlear filters and linearization of the cochlear input-output function. High-rate AM is processed in cochlear filters with high characteristic frequencies (CFs). It is possible that broader filters and more linear cochlear response at high CFs facilitated 100-Hz AM detection in older listeners. According to this reasoning, the middle and older groups should have also performed better than the young group for the two higher modulation rates, which was not the case. Since the modulated noise was presented at an overall level of 70 dB SPL, the advantage from OHC loss in older adults might have been offset by lower stimulus level at the output of their high-CF cochlear filters (due to hearing loss). In fact, for some listeners in the middle and older groups, the noise components within high-CF filters may have fallen close or even below hearing thresholds.

To further investigate the relative contributions of age and high-frequency hearing loss to the detection of high-rate AM, Pearson’s product correlations were calculated, for each modulation rate separately, using AM detection thresholds pooled for all the listeners from the three age groups as the dependent variable and age and HFAVG as independent variables. We then performed partial correlations, one with age after controlling for the HFAVG, and another with HFAVG after controlling for age. The results are shown in Table 2. Correlation of AM-detection thresholds with age was only significant for the 100-Hz rate but it became non-significant after regressing out HFAVG. However, correlation between AM-detection thresholds and HFAVG was significant for two modulation rates, 100 and 300 Hz, and remained significant after age was regressed out. The results of the correlation analyses support our argument that high-frequency hearing loss was the main contributor to the improvement in AM detection in older listeners for modulation rates processed in cochlear filters producing audible output to the noise. Our finding is consistent with the improvement of AM-detection threshold with increasing hearing threshold at the carrier frequency observed by Prendergast et al. (2019) for a lower modulation rate (25 Hz).

**Table 2.** Person’s product correlation coefficients for relationships between AM-detection thresholds and age, AM-detection thresholds and average hearing thresholds across 2 – 8 kHz (HFAVG). Correlation coefficients are shown for the three modulation rates used, listed in column 1. Two rightmost columns show partial correlations with the controlled variable in parentheses. Significant correlations are shown in bold font (** p < 0.01 and * p < 0.05).

| AM rate [Hz] | Age | HFAVG | Age (HFAVG) | HFAVG (Age) |
| --- | --- | --- | --- | --- |
| 100 | **-0.34\*\*** | **-0.40\*\*** | -0.15 | **-0.26\*** |
| 300 | -0.03 | **-0.27\*** | 0.16 | **-0.31\*** |
| 600 | -0.04 | -0.19 | 0.08 | -0.20 |

The lack of adverse age effects on AM detection is inconsistent with electrophysiological data reported by Purcell et al. (2004), which showed degraded coding of AM for rates above 100 Hz in older compared to younger listeners. The age within the older group in that study extended up to 78 yrs (compared to 66 yrs in this study). It may be that age-related deficits in the processing of high-rate AM occur at older ages than those of our participants. Overall, we found no evidence of degraded AM processing with age, as measured by AM-detection thresholds for high modulation rates (100-600 Hz) when controlling for listeners’ hearing sensitivity.

#### 2.3.2 Gap detection for tones in noise

A subset of 53 of the 61 participants completed a gap-detection task for tones presented in noise. The average gap detection thresholds converted to milliseconds, for the three age groups are plotted in Fig. 3. The left and right sets of bars show the thresholds for detecting gaps in the 2- and 4-kHz kHz tones, respectively. As evident in the figure, listeners in all three age groups were able to detect shorter gaps at 4 kHz than at 2 kHz.

**Fig. 3.**
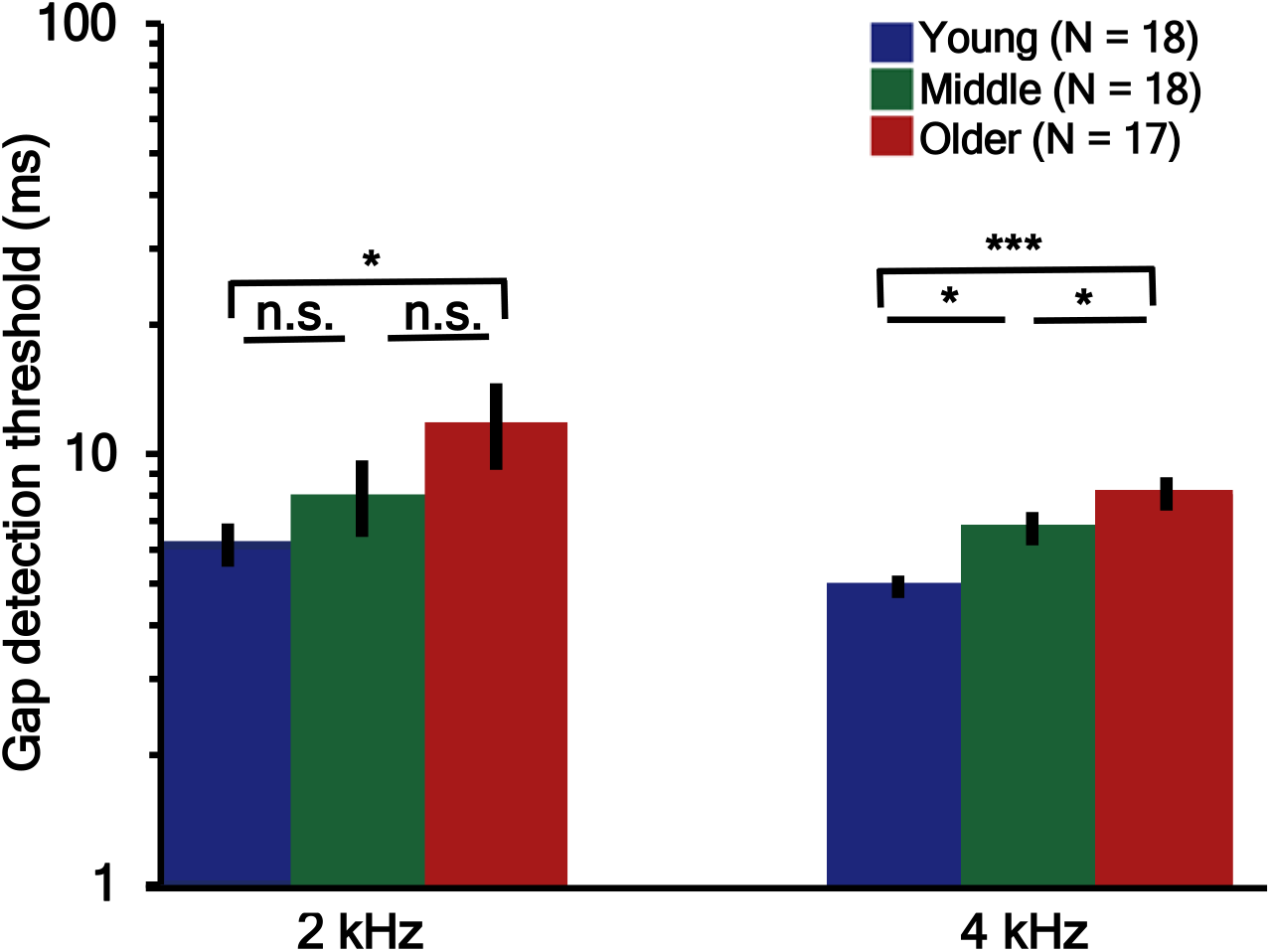
Gap-detection thresholds for 2- and 4-kHz tones presented in one-octave noise bands, for three age groups. The error bars indicate ± 1 SE. Asterisks denote significant differences (*** p < 0.001, and * p < 0.05).

A mixed-design ANOVA performed on log-transformed gap-detection thresholds with the frequency of the tonal marker as the within-subjects factor, the age group as the between-subjects factor, and the HFAVG (defined above) as a covariate, showed a significant effect of marker frequency [F(1, 49) = 21.32, p < 0.001]. There was also a significant effect of age group [F(2, 49) = 5.64, p = 0.006], but no significant effect of HFAVG [F(1, 49) = 0.19, p = 0.67]. In addition, we found no significant interactions between the marker frequency and age group [F(2, 49) = 0.21, p = 0.816] or between marker frequency and HFAVG [F(1, 49) = 1.35, p = 0.251]. Because of that, post hoc pairwise comparisons were performed using thresholds pooled across the two marker frequencies. The comparisons showed a significant difference in sensitivity to temporal gaps between the young and older groups (p = 0.005), but not between the young and middle (p = 0.223) or middle and older (p = 0.188) groups.

Treating listeners’ age as a continuous variable, Pearson’s correlations were performed separately for each tone frequency, with gap-detection thresholds as the dependent variable and age and HFAVG as independent variables. As shown in Table 3, neither age nor HFAVG were significantly correlated with listeners’ ability to detect gaps in the 2-kHz tone, although correlation with age approached significance with and without controlling for HFAVG (p = 0.051 and p = 0.052, respectively). At 4 kHz, gap-detection thresholds were significantly correlated with age and the correlation remained very strong after controlling for HFAVG. There was also a significant correlation with HFAVG at 4 kHz but it became non-significant after controlling for age. In sum, the results of the correlational analyses show that age but not high-frequency hearing sensitivity contributed to a decline in the ability to detect temporal gaps in a 4-kHz tone presented in noise.

**Table 3.**
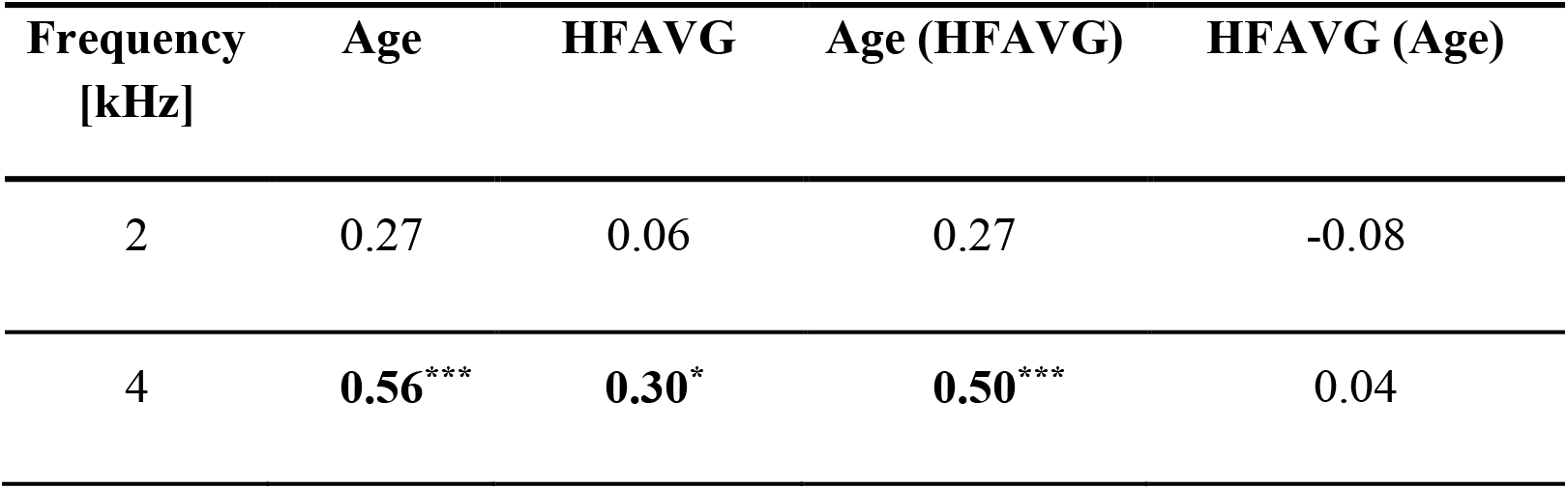
Pearson’s product correlation coefficients for relationships between gap-detection thresholds and age as well as HHAVG (2-8 kHz). Correlations were performed separately for a 2- and 4-kHz tonal markers of a gap. Two rightmost columns show partial correlations with the controlled variable listed in the parentheses. Significant correlations are shown in bold font (*** p < 0.001 and * p < 0.05).

Overall, the data are consistent with a large number of studies that reported increased gap detection thresholds with advancing age (Grose et al., 2006; Humes et al., 2009; Humes et al., 2010; Kumar et al., 2011; Pichora-Fuller et al., 2006). It has been found that the gap detection performance deteriorates at middle age (> 40 yr) compared with that for young adults (Grose et al., 2006; Kumar et al., 2011). Our data show this pattern for the 4-kHz tone presented in noise and a trend consistent with it for the 2-kHz tone. In contrast, a recent study by Schoof et al. (2014) did not find a significant difference in listeners’ ability to detect a gap in wideband noise between older and younger groups tested in that study. It may be that a smaller sample size combined with large variability in the data contributed to the null age effect in that study. The statistical analyses show that the decline in the ability to detect brief gaps in stimuli presented at suprathreshold levels in noise with advancing age appears independent of hearing loss that is often concomitant with aging (e.g., Humes et al., 2010).

#### 2.3.3 Interaural time difference in stimulus envelope

Thresholds for detecting envITDs for two carrier frequencies (2 and 4 kHz) and two modulation rates (40 and 100 Hz) were obtained for a subset of 27 listeners. The listeners completed the task in quiet and in noise. The log-transformed envITDs at threshold averaged within each of the three age groups and converted to linear units (μs) are shown in Fig. 4. The left and right panels show data for the 40- and 100-Hz AM, respectively. In each panel, the left and right set of bars show the thresholds measured for the 2- and 4-kHz carrier, respectively, in quiet and in noise.

**Fig. 4.**
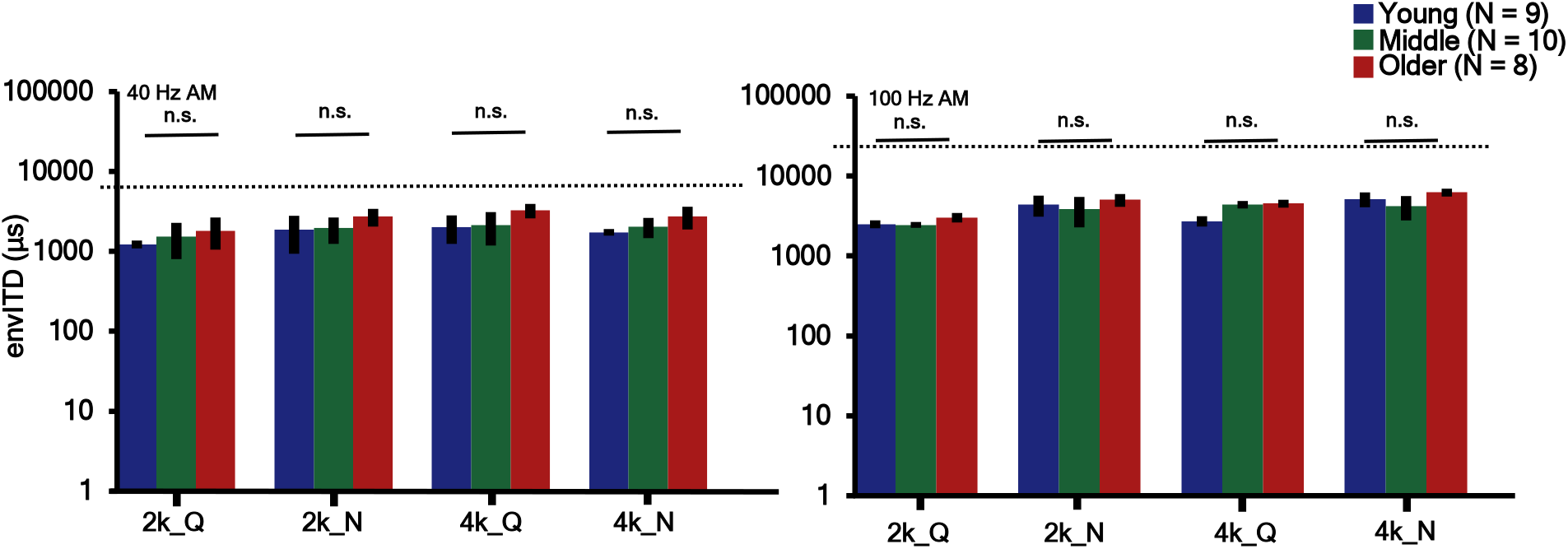
Thresholds for detecting interaural time difference in envelope (envITD), for three age groups. The error bars indicate ± 1 SE. Data for the 40- and 100-Hz AM are shown in the left and right panel, respectively. Thresholds were measured for 2- and 4-kHz carriers in quiet (Q) and in one-octave noise (N). The horizontal dotted lines indicate envITD corresponding to an interaural envelope phase difference of 180 deg, for the 40-Hz (left panel) and 100-Hz (right panel) AM rates.

A mixed-design ANOVA was performed on log-transformed envITDs for the two modulation rates. The modulation rates were considered separately since comparable phase shifts for the 40- and 100-Hz rate correspond to different envITDs. In both cases, carrier frequency and condition (quiet\in noise) were used as within-subjects factors, age group was used as the between-subjects factor, and HFAVG was used as a covariate. For the 40-Hz AM rate, there was no significant effect of carrier frequency [F(1, 23) = 0.726, p =0.403] and no significant effect of the presence/absence of noise [F(1, 23) = 1.31, p = 0.299]. There was also no significant effect of age group [F(2, 23 = 1.88, p = 0.175] or HFAVG [F(1, 23) = 0.451, p = 0.508] on envITD thresholds. None of the two- and three-way interactions were statistically significant (p > 0.05 in all cases). For the 100-Hz AM rate, the effect of carrier frequency was significant [F(1, 23) = 15.09, p = 0.001] indicating significantly worse performance for the 4-kHz than 2-kHz carrier. The effect of condition (quiet/in noise) was also significant [F(1, 23) = 5.61, p = 0.027] indicating that the presence of noise had an adverse effect on envITD thresholds. However, there was no significant effect of age group [F(2, 23) = 0.87, p = 0.432], no significant effect of HFAVG [F(1, 23) = 0.014, p = 0.907], and no significant two- and three-way interactions of the two factors with carrier frequency and condition (p > 0.05).

For participants pooled together from the three age groups, there were no significant correlations between age and envITD thresholds, between HFAVG and envITD thresholds, and no significant partial correlations with either of the two variables with the other controlled for (p > 0.05). This was true for all the conditions tested (two carrier frequencies, two modulation rates, in quiet and in noise).

The absence of age effects on sensitivity to envITDs was also reported by Prendergast et al. (2019), despite the fact that Prendergast et al. (2019) used a higher modulation rate (255 Hz), for which previous studies have found deficits in monaural processing of AM (Kumar et al., 2011; Purcell et al., 2004). However, the lack of significant correlation with hearing sensitivity in the frequency range corresponding to carrier frequencies contrasts with the significant contribution of hearing sensitivity to envITDs reported by Prendergast et al. (2019). One reason for the discrepant findings may be a smaller sample size in our study. The correlations reported by Prendergast et al. (2019) showed that hearing sensitivity predicted only about 11% of variance in envIPD thresholds, a small effect that perhaps could not be detected with a relatively small sample used here, given the inherent variability in envITD thresholds both within and across listeners.

Still, some earlier studies reported a decline in the binaural envelope processing with age even as the differences in hearing sensitivity were accounted for. King et al. (2014) reported a significant correlation between thresholds for detecting envITDs and age for low-frequency carriers (250 and 500 Hz) modulated at a relatively low rate (20 Hz). The upper age limit of their listeners was 83 and the significant correlations were mainly due to poor performance of those above 65 yrs of age (near the upper limit of age for the older group in this study). It is possible that deficits in envITDs emerge at a later age above the age range of participants in our and Prendergast et al. (2019) studies.

## 3 Electrophysiological envelope following responses

### 3.1 Participants

Electrophysiological EFRs were measured in 45 (33 female) of the 61 initially recruited participants. The remaining participants were unavailable for testing due to their time constraints. Most of the participants completed at least two of the psychophysical tasks described above in Section 2.

### 3.2 Stimuli and procedure

EFRs were measured in response to a series of 437.52-ms long tones modulated in amplitude at a rate of 91.42 Hz. Of the 45 participants, 21 participants (6 young, 8 middle, and 7 older) with normal hearing at all audiometric frequencies were tested using a carrier frequency of 4 kHz. Because 8 older participants had some clinical hearing loss at 4 kHz, the remaining 24 participants (8 young, 8 middle, and 8 older) were tested using a carrier frequency of 2 kHz. The AM tones were presented in quiet in one condition and in a notched noise in another condition. The EFRs were measured for two modulation depths, −8 and 0 dB [20log(*m*)], for AM tones presented in quiet. For tones in noise, the EFRs were measured for three modulation depths, −8, −4, and 0 dB. The noise had a bandwidth extending from 0.1 to 10 kHz with a spectral notch centered on the carrier frequency. The notch width was 0.2 *x* carrier frequency. In each condition, the AM tones were presented 1000 times with an average inter-stimulus-interval of 550 ms with a small time jitter distributed uniformly over a 20-ms interval around the mean. The stimuli were presented with positive polarity in half of the trials and negative polarity in the other half. The order of the five test conditions (quiet *x* 2 modulation depths, in noise *x* 3 modulation depths) was randomized across participants. The AM tones were presented at 70 dB SPL, and the notched noise had an overall level of 60 dB SPL.

Participants were seated in an electrically shielded double-walled sound-attenuating booth and neural responses were recorded using a 64 channel EEG system (Biosemi Active II system). Participants wore a fitted cap (Easy Cap, Falk Minow Services) containing 64 silver-chloride scalp electrodes. In addition, a reference electrode was placed on the mastoid of the test ear, and two additional ocular electrodes were used to record eye movement and eyeblink activity. The voltage DC offset was maintained below ± 20 mV for all electrodes. The neural activity in each channel was recorded at a sampling rate of 4096 Hz. A desktop computer was used to present and trigger the stimuli on the Biosemi software and store the data. The stimuli were generated with a sampling rate of 44.1 kHz and were routed to a Tucker-Davis Technologies System 3 (Gainsville, FL) for monaural presentation via an ER-1 insert phone with a foam ear tip (Etymotic Research, Elk Grove Village, IL). To divert attention from the stimuli and avoid boredom during the measurements, participants watched a silent captioned movie of their choice.

The pre-processing and averaging of the EEG recordings was performed using the EEGLAB toolbox (Delorme et al., 2004). The raw waveforms were first downsampled to 1024 Hz, re-referenced to the test-ear mastoid, and band-pass filtered into a range from 1 to 350 Hz using a 4th-order Butterwoth filter. A zero phase shift was achieved by using Matlab’s *filtfilt* function. The continuous EEG time waveforms were divided into epochs, from −100 ms before stimulus onset to 500 ms post onset. The 100-ms pre-stimulus interval was used for baseline correction. Independent Component Analysis (ICA) was conducted to isolate and remove artifacts related to eye movements and eyeblinks (Jung et al., 2000). Discrete Fourier transforms of the pre-processed EEG waveforms from individual trials were performed to obtain the phase spectrum. For each electrode, the phase locking value (PLV) to the AM in the stimulus was calculated by averaging the phases at each frequency across 800 randomly selected samples from individual trials (Zhu et al., 2013). A bootstrapping technique, as described by Zhu et al. (2013), but for 800 trials, was used to estimate the average PLV and the noise floor. Based on this analysis, a PLV was considered significant when it exceeded a value of 0.045. A subset of 35 electrodes that yielded the largest PLVs in the test ear hemisphere was selected for averaging to calculate the average PLV that was used as the final estimate for each subject and each condition. These PLV estimates were used for statistical comparisons performed to test for effects of age and HFAVG on neural responses to AM.

### 3.3 Results and discussion

All the estimated PLVs were above the limit corresponding to the 95th percentile of the noise floor distribution and thus all were included in statistical analyses. The PLVs averaged separately across participants who completed the experiment for the 2- and 4-kHz carriers are shown in the left and right panels of Fig. 5, respectively. The different color bars show the PLVs for different age groups. As evidenced by the bar plot, the presence of the notched noise resulted in substantially reduced PLVs compared with those observed for the AM tones presented in quiet.

**Fig. 5.**
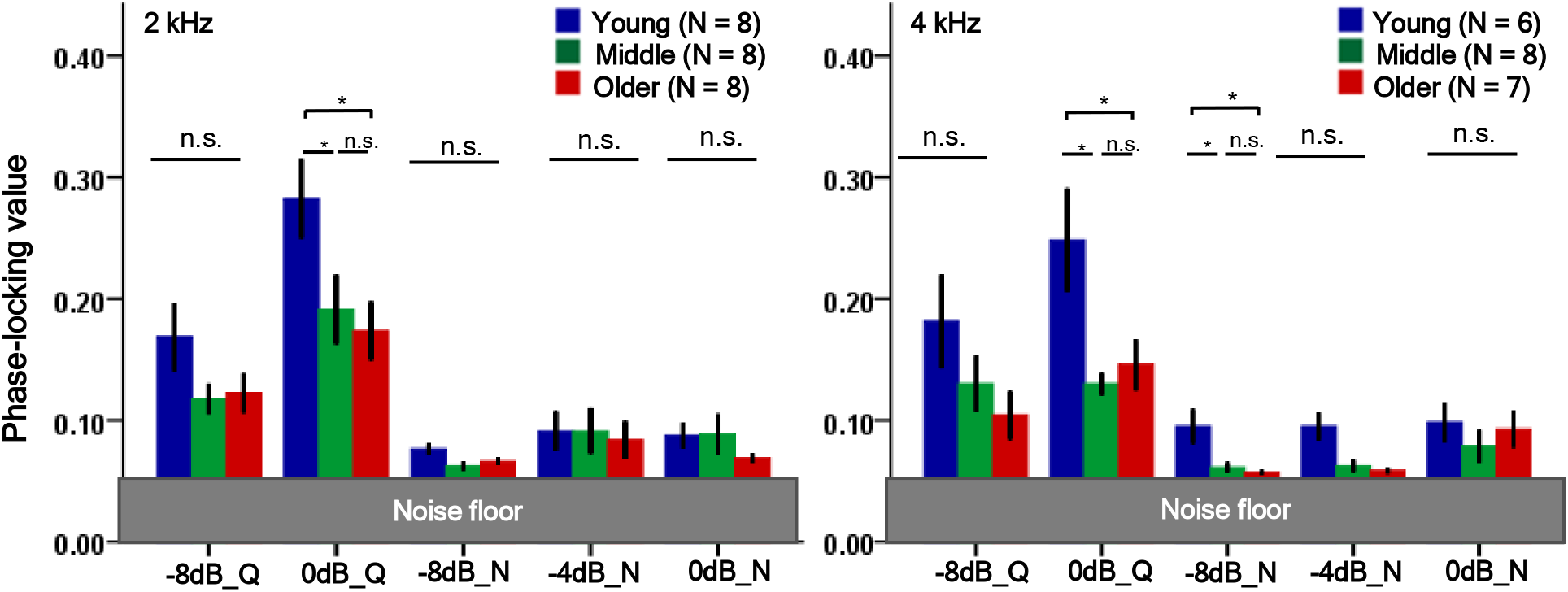
EFR phase-locking values for a 91.4-Hz AM in a 2-kHz carrier (left panel) and a 4-kHz carrier (right panel). The error bars indicate ± 1 SE of the mean. The EFRs were measured for two modulation depths, −8 and 0 dB, in quiet (Q), and for three modulation depths, −8, −4, and 0 dB, in noise (N). The asterisks indicate significant differences (* p < 0.05).

Mixed-design ANOVAs were performed separately on PLVs measured using the 2- and 4-kHz carriers. The five test conditions (quiet\noise *x* 2\3 modulation depths) were used as within-subjects factors, the age group was used as the between-subjects factor, and HFAVG (2 to 8 kHz) was used as a covariate. Because Mauchly’s test showed that the assumption of sphericity was violated for both carrier frequencies, a Greenhouse-Geisser correction was applied in both cases. For the 2-kHz carrier, the ANOVA showed a significant effect of condition [F(2.23, 80) = 16.92, p < 0.001], significant effect of age group [F(2, 20) = 3.91, p = 0.037], but no significant effect of HFAVG [F(1, 20) = 1.32, p = 0.265]. There was no significant interaction between condition and age group [F(4.47, 80) = 1.90, p = 0.121] or between condition and HFAVG [F(2.23, 80) = 0.64, p = 0.548]. The Games-Howell test of multiple pairwise comparisons showed a significant difference between the young and older groups for the 0-dB modulation depth in quiet (p = 0.048) but the significance would not survive correction for multiple comparisons. In all other conditions, none of the pairwise group comparisons returned a significant effect. For the 4-kHz carrier, the ANOVA showed a significant effect of condition [F(2.36, 68) = 8.66, p < 0.001], a significant effect of age group [F(2,17) = 6.28, p = 0.009], but no significant effect of HFAVG [F(1, 17) = 0.008, p = 0.931]. There was no significant interaction between condition and age group [F(4.71, 68) = 1.76, p = 0.146] or between condition and HFAVG [F(2.36, 68) = 0.77, p = 0.488]. The Games-Howell test showed significant differences between the young and middle groups (p = 0.01) and between the young and older groups (p = 0.031) for the 0-dB AM depth in quiet, and between the young and middle groups (p = 0.017) and the young and older groups (p = 0.009) for the −8-dB AM in noise. Only the latter effect would survive correction for multiple comparisons. No other pairwise differences were significant (p > 0.05).

Correlations were performed on EFR PLVs pooled across all age groups and the two carrier frequencies, separately for each of the five experimental conditions (quiet\noise *x* 2\3 modulation depths), with the PLV as the dependent variable and age and HFAVG as independent variables. None of the correlations and none of the partial correlations with either dependent variable was statistically significant after controlling for the other (p > 0.05).

One reason for the lack of age effects on EFRs could be that the modulation rate used in this study was not sufficiently high to reveal deficits in older individuals (Purcell et al., 2004). Other studies using similar (~100 Hz) and lower modulation depths also reported no effects of age (Bharadwaj et al., 2015; Prendergast et al., 2019) although Bharadwaj et al. (2015) only tested listeners younger than 40 yrs. Zhu et al. (2013) and Bharadwaj et al. (2015) found a significant relationship between the slope of a function relating EFR spectral magnitudes to the modulation depth and listeners’ noise exposure. They argued that the slope-based measure was a better summary measure than the EFR magnitude itself as it provided normalization that removed influences of non-auditory factors on EEG recordings (Mitchell et al., 1989), thus facilitating between-group comparisons of AM processing. In both these studies, the slopes were steeper for individuals whose data showed patterns consistent with the presence of cochlear synaptopathy. In an apparent contrast, Garrett et al. (2019) observed shallower EFR slopes in older listeners than in their young normal-hearing sample. Garrett et al. (2019) suggested that high-frequency hearing loss in the older listeners may have offset the effect expected from cochlear synaptopathy. However, the findings of the steeper EFR slopes were also not replicated in a large scale study by Prendergast et al. (2017a) who found no relationship between EFR slopes and noise exposure. In this study, the data in quiet were collected for only two modulation depths, yielding highly variable slope estimates. However, we fitted straight lines to PLVs observed for the three modulation depths for the AM tones presented in the notched noise. ANOVAs conducted separately on the slopes for the 2- and 4-kHz carriers showed no effect of age group [F(2,23) = 0.44, p = 0.653, for the 2-Hz carrier, and F(2, 20) = 0.74, p = 0.491, for the 4-kHz carrier, and no effect of HFAVG [F(2, 23) = 3.11, p =0.093 for the 2-kHz carrier, and F(2, 20) = 0.52, p = 0.481 for the 4-kHz carrier]. Overall, except for isolated conditions we found no effect of age and HFAVG on either the PLVs or the slopes of functions relating PLVs to the modulation depth.

## 4 Speech perception and spatial release from masking

### 4.1 Participants

Fifty-eight listeners (36 females) completed the speech experiment. The number included 27 listeners who performed all the experiments in this study. The rest performed at least one other experiment.

### 4.2 Stimuli and procedure

The speech stimuli were sentences from the Harvard Institute of Electrical and Electronics Engineers (IEEE, 1969) corpus spoken by a female with a voice with a pitch (F0) of ~ 180 Hz. The background speech consisted of a mixture of two female voices that had either the same pitch as the target (same-pitch condition) or were processed using Praat software (Boersma et al., 2010) to yield one background voice pitch that was three semitones below, and the other three semitones above that of the target (different-pitch condition). The spatial location of the target and background speakers was simulated by applying non-individualized head-related transfer functions to the stimuli. In one condition, the background speakers were colocated with the apparent target location corresponding to a 0-deg azimuth. In another condition, the target speech was at a 0-deg azimuth while the background speakers had different spatial locations, one corresponding to the +15-deg and the other to −15-deg azimuth. To make the task more challenging, the speech stimuli were also convolved with an impulse response of a reverberant room (T60 = 1.07s). The four different conditions, 2 target/maskers voice pitch (same/different) configurations *x* 2 speaker locations (colocated/non-colocated) were presented in a randomized order for each listener. In all conditions, the target speech was presented at 75 dB SPL. In the same-pitch conditions, the signal-to-noise ratio (SNR) was 2 dB, while in the different-pitch conditions, the SNR was lowered to 0 dB to prevent ceiling performance. For older listeners who had high-frequency hearing thresholds above 20 dB HL, the stimuli were amplified in the region of hearing loss using a half-gain rule.

In addition, the four conditions described above were repeated using speech stimuli from the same corpus that were lowpass filtered to limit their spectra to the frequency range over which all the listeners had normal hearing sensitivity by clinical standards. A 4th-order Butterworth lowpass filter with a cutoff frequency of 2 kHz was used to limit the spectrum. The filtering was performed using Matlab’s *filtfilt* function to remove phase shifts across frequency. Spread of excitation into high-frequency regions of the cochlea was masked by presenting the lowpass-filtered speech with a highpass-filtered Gaussian noise. A 4th-order Butterworth filter with a cutoff frequency of 2200 Hz was used to create the masking noise. The level of the noise was 10 dB below that of the target speech. For the lowpass-filtered speech, the SNR was set to 6 dB in the same-pitch conditions and to 4 dB in the different-pitch conditions. The two-talker background babble was created by concatenating sentences spoken by the maskers from lists that were not used for the target speech in any test conditions for a given listener. The masker waveforms were summed and segmented so that the masking speech was longer than the target by at least 1 second (the masker duration was set relative to the longest sentence in the corpus). In each trial, the target started after a 0.8 sec delay from the onset of the two-talker babble. The highpass noise in the lowpass speech condition was 2 sec longer than the two-talker masker. The two-talker masker with the target started 0.8 sec after the noise onset.

The experiment began with practice during which one list was presented for each test condition using full speech spectrum. None of the lists used during the practice was repeated during the testing. Two lists of 10 sentences (a total of 20 sentences) were used for each test condition. After each sentence, listeners were prompted to type the sentence they heard. If they missed parts of the sentence, they were asked to type any number of words they heard and they were encouraged to make the best guess even if they were not sure about the exact words. No feedback was provided during training or data collection. Five keywords in each sentence were used to obtain a score. Words that were clearly misspelled (rather than not heard) were counted as correct responses. The proportion of correctly recalled keywords across two lists was used as the final score for each test condition.

During the testing, participants were seated in a single-walled sound-attenuating booth. The stimuli were played in Matlab (Mathworks, Natick, MA) using an E22 soundcard (LynxStudio, Costa Mesa, CA) with a sampling rate of 44.1 kHz and were delivered to both ears via Sennheiser HD650 headphones (Sennheiser, Old Lyme, CT).

### 4.3 Results and discussion

The proportion correct scores from each condition were converted to rationalized arcsine units (RAUs; Studebaker, 1985). Fig. 6 shows the mean RAU scores for the same- and different-pitch condition in the upper and lower row of panels, respectively. The plots in the left column show the scores for the full-spectrum speech and those in the right column show the scores for the lowpass-filtered speech. The unfilled and hatched bars show data for the non-colocated and colocated speakers, respectively. The scores for the corresponding colocated and non-colocated conditions (within each panel) were obtained using the same SNRs, and the scores in different panels were obtained using different SNRs, as indicated in Section 4.2, and thus cannot be directly compared.

**Fig. 6.**
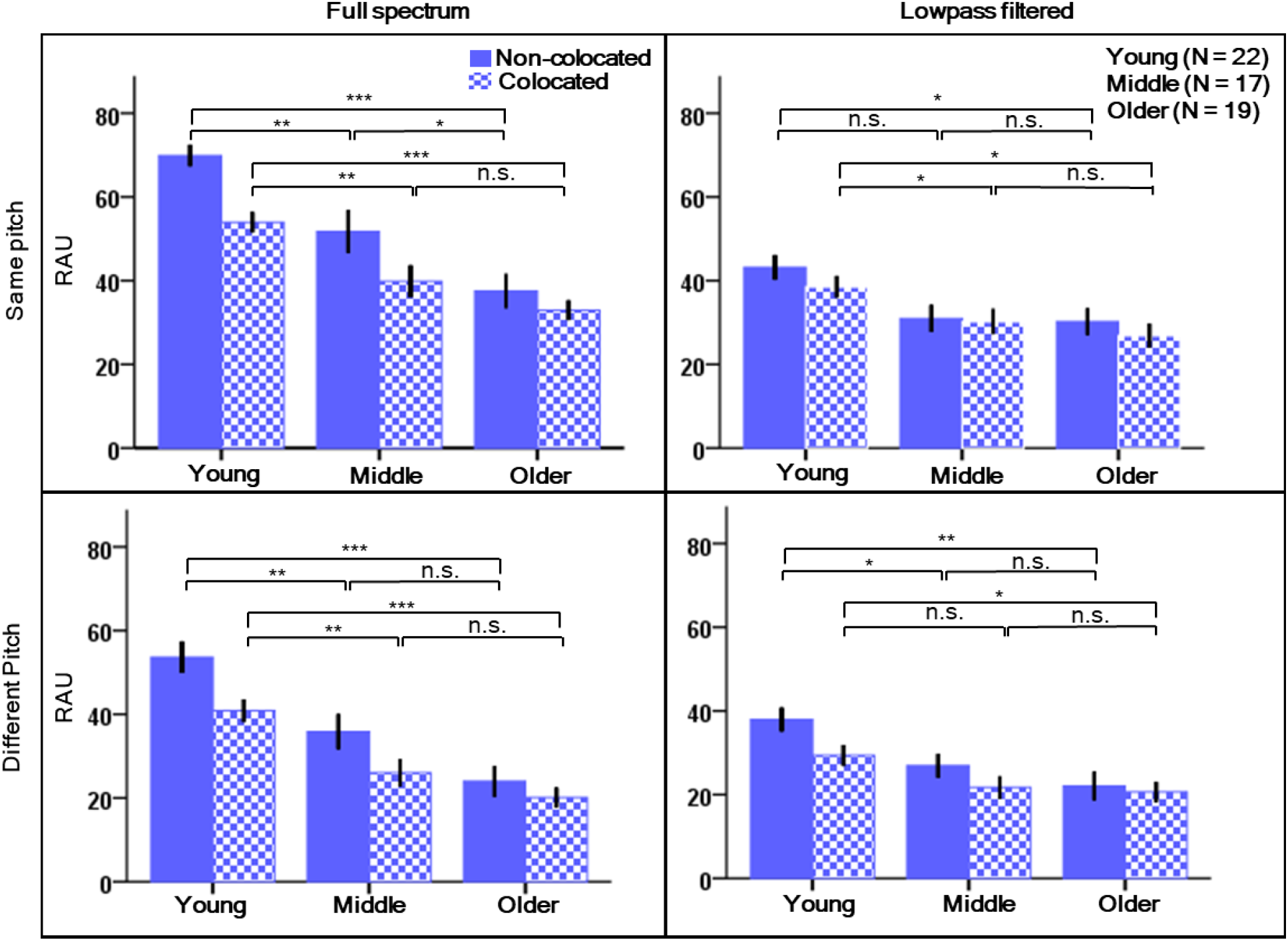
Speech scores (in RAUs) for target speech in two-talker babble. Unfilled bars show the scores for the target at 0-deg azimuth and the competing speakers at +15-deg and −15-deg azimuths. The error bars indicate ± 1 SE of the mean. The upper panels show data for the target and masker speakers with the same voice pitch, and the lower panels show the data for one masker with voice pitch three semitones below, and the other three semitones above that of the target. The left and right column of panels show data for the full-spectrum and the lowpass-filtered speech, respectively. Asterisks denote significant differences (*** p < 0.001, ** p < 0.01, and * p < 0.05).

Four mixed-design ANOVAs were performed on the RAU scores separately for the four conditions (2 voice pitch x 2 speech spectrum), with relative spatial location of the target and maskers (colocated\non-colocated conditions) as the within-subjects factor, age group as the between-subjects factor, and average hearing threshold as a covariate. For the full-spectrum speech, the average hearing threshold was calculated across frequencies 0.25 to 8 kHz (ALLAVG) because there were significant differences in hearing thresholds between the three age groups across the entire range of audiometric frequencies (see Fig. 1). For lowpass-filtered speech, the average hearing threshold was calculated for octave frequencies from 0.25 to 2 kHz (LFAVG) because speech information was limited to this frequency range.

The ANOVA for the same-pitch full-spectrum condition showed a significant effect of spatial condition [F(1, 54) = 14.17, p < 0.001], a significant effect of age group [F(2, 54) = 5.34, p = 0.008], and a significant effect of ALLAVG [F(1, 54) = 4.67, p = 0.035]. There was no significant interaction between condition and age group [F(2, 54) = 1.21, p = 0.307] or between condition and ALLAVG [F(1, 54) = 1.06, p = 0.308]. Post-hoc pairwise comparisons showed that in the colocated condition, the young group performed significantly better than the middle-age group (p = 0.003) and the older group (p <0.001), but there was no significant difference in speech scores between the middle and older groups (p = 0.239). Performance in the non-colocated condition declined progressively with age (young vs. middle, p = 0.006; young vs. older, p < 0.001; middle vs. older, p = 0.04), indicating a decline of the ability to use a small spatial separation (± 15 deg) between the target and masker speakers to improve performance.

The ANOVA for the different-pitch condition showed no significant difference between the scores for the colocated and non-colocated target and maskers [F(1, 54) = 3.34, p = 0.073]. This result suggests that different voice pitch was a sufficient cue to segregate the target and maskers in the colocated condition and that spatial segregation did not provide an additional benefit for speech understanding. There was, however, a significant effect of age group [F(2, 54) = 6.06, p = 0.004], but no significant interaction between the condition and age group [F(2, 54) = 1.77, p = 0.181] indicating an overall worse performance of older than younger individuals in both colocated and non-colocated conditions. Post-hoc pairwise comparisons showed significant differences between the young and middle groups (p = 0.001) and the young and older groups (p < 0.001), but not between the middle and older groups (p = 0.328). Similarly, in the non-colocated condition, there was a significant difference in performance between the young and middle (p = 0.006) and young and older (p < 0.001), but not middle and older (p = 0.103) groups. The ANOVA showed no significant effect of ALLAVG [F(1, 54) = 2.93, p = 0.093] and no interaction between spatial condition and ALLAVG.

The ANOVA performed on RAU scores for the same-pitch lowpass-filtered condition showed no significant effect of spatial condition [F(1, 54) = 0.41, p = 0.526] indicating that spatial segregation cues were insufficient in the low-frequency speech spectrum to improve speech recognition in the non-colocated compared to colocated condition. There was a significant effect of age group [F(2, 54) = 3.48, p = 0.038], but no significant interaction between spatial condition and age group [F(2, 54) = 0.68, p = 0.513] indicating that in both spatial conditions, older participants performed worse than younger participants. Post hoc pairwise comparisons showed that in the colocated condition, the young group performed significantly better than the middle (p = 0.021) and the older (p = 0.01) groups, but there was no significant difference in performance between the middle and older groups (p = 0. 985). In the non-colocated condition, there was a significant difference in performance between the young and older groups (p = 0.01) but not between the young and middle (p = 0.103) or between middle and older (p = 0.693) groups. For the same-pitch lowpass-filtered speech, the ANOVA also showed a significant effect of LFAVG [F(1, 54) = 5.67, p = 0.021], but no interaction between spatial condition and LFAVG [F(1, 54) = 0.47, p = 0.497].

For the different-pitch lowpass-filtered speech condition, the ANOVA showed no significant effect of spatial condition [F(1, 54) = 1.80, p = 0.185] but there was a significant effect of age group [F(2, 54) = 4.54, p = 0.015] and a significant interaction between condition and age group [F(2, 54) = 3.69, p = 0.032]. Post hoc pairwise comparisons revealed that for the colocated condition there was a significant difference in performance between the young and older groups (p = 0.041) but not between the young and middle (p = 0.098) or middle and older (p = 0.952) groups. In the non-colocated condition, the young group performed significantly better than both the middle (p = 0.037) and older (p = 0.001) groups but there was no significant difference between the middle and older groups (p = 0.534). The ANOVA showed that LFAVG was not a significant factor [F(1, 54) = 0.58, p = 0.451] and there was no significant interaction between LFAVG and spatial condition for the low-pass filtered speech with different target/masker voice pitches [F(1, 54) = 1.85, p =0.18].

Overall, the pairwise comparisons showed that for full-spectrum speech, significant decline in speech perception is already observed in middle-aged individuals compared to the young adults and this is true when the target and maskers are colocated or non-colocated with relatively small spatial separation. The age effects on perception of speech in the colocated condition in this study appear in contrast with the lack of significant effects of age in speech-on-speech masking measured using diotic presentation in the study by Prendergast et al. (2019). Differences in experimental procedures may have contributed to the different outcomes. Prendergast et al. (2019) used a closed-set speech corpus (coordinated response measure) for both the target and the two-talker masker while IEEE sentences were used in this study. Another difference was that Prendergast et al. (2019) measured SNRs needed for performance corresponding to 25% correct word recognition. In this study SNR = 0 dB was used in the comparable (diotically presented full-spectrum speech) condition. For a low-pass filtered speech, our finding of significant age effects on performance in the colocated condition is consistent with that reported by Leger et al. (2014) for perception of vowel-consonant-vowel syllables in the presence of a single interfering talker.

The RAU scores were used to calculate SRM separately for each voice-pitch and speech-spectrum condition by subtracting the score for the colocated target and maskers from the score for the non-colocated target and maskers. The magnitudes of SRM for the full-spectrum and the lowpass-filtered speech are shown in the left and right panels of Fig. 7, respectively. In each plot, the left three bars show SRM for the same-pitch, and the right set of bars, for the different-pitch conditions.

**Fig. 7.**
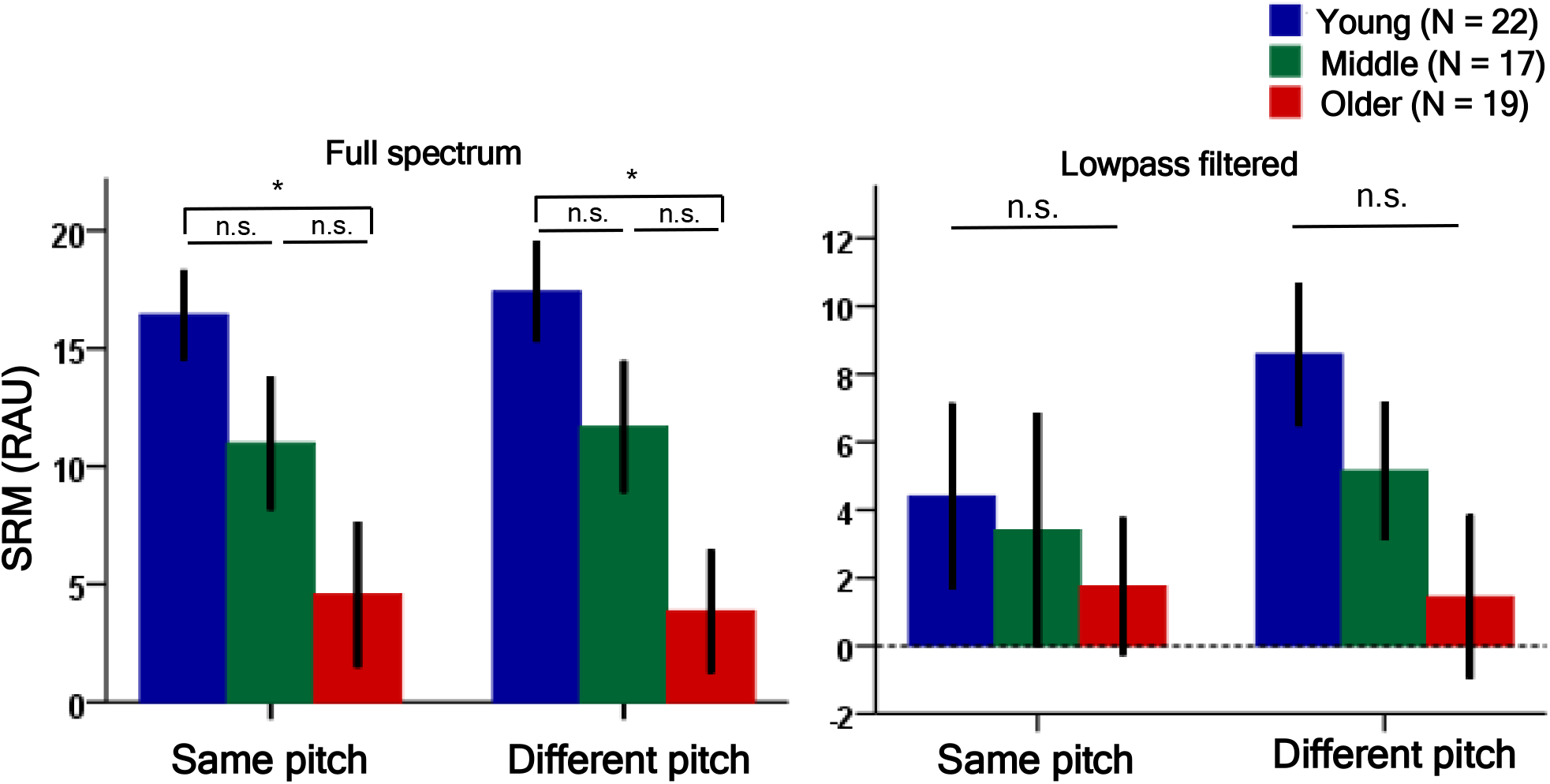
Spatial release from masking, for three age groups, for full-spectrum speech (left panel) and for speech lowpass-filtered at 2 kHz (right panel). In each panel, the left and right set of bars show SRM for the same-pitch and different-pitch conditions, respectively. The error bars indicate ± 1 SE of the mean. Asterisks denote significant differences (* p < 0.05).

Although age group was a significant factor for speech understanding in all four speech conditions (2 voice pitch *x* 2 speech spectrum), the SRM, which is a differential measure reflecting benefits of spatial segregation, was not always affected by age. To further investigate the effects of age and hearing sensitivity on SRM, we performed correlations separately for each voice-pitch and speech-spectrum conditions, for participants pooled across the three age groups. The scatterplots in Fig. 8 show the individual magnitudes of SRM for the full-spectrum speech (left panel) and the lowpass-filtered speech (right panel). In both panels, the grey and red circles show the data for the same- and different-pitch conditions, respectively.

**Fig. 8.**
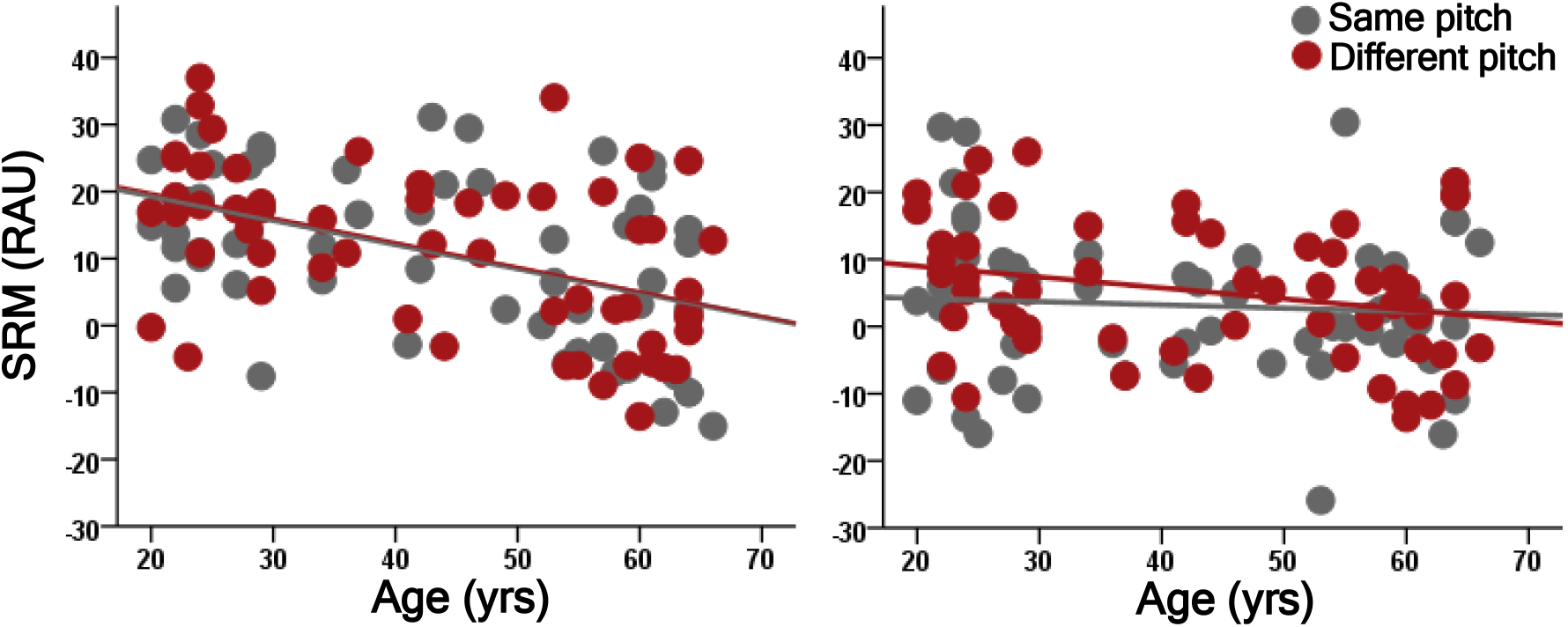
Scatterplots in the left and right panels show individual SRM plotted as a function of participants’ age for the full-spectrum and lowpass-filtered speech, respectively. Grey and red symbols show data for the same-pitch and different-pitch conditions, respectively. The lines represent linear regression fit to the data (see Pearson’s product correlations coefficients in Table 4).

The results of the correlational analyses are reported in Table 4. For the full-spectrum speech, the outcomes were similar for the same-pitch and different-pitch conditions. In both cases, there was a significant correlation between SRM and age and the correlation remained significant after partialling out ALLAVG. There was also a significant correlation with ALLAVG but the correlation became non-significant when age was controlled for. For the lowpass-filtered speech, neither age nor LFAVG were significantly correlated with the size of SRM.

**Table 4.** Correlations between spatial release from masking (SRM) and age, and between SRM and average hearing threshold (AVG_HT). For the full-spectrum speech, AVG_HT was calculated using thresholds in dB SPL over the frequency range from 0.25 to 8 kHz, while for the lowpass-filtered speech AVG_HT was calculated using thresholds from 0.25 to 2 kHz. Variables listed in parentheses were controlled for in partial correlations shown in the two rightmost columns. Significant correlations are shown in bold font (*** p < 0.001, ** p < 0.01, and * p < 0.05).

| SRM | Age | AVG_HT | Age<br>(AVG_HT) | AVG_HT<br>(Age) |
| --- | --- | --- | --- | --- |
| Same pitch (full spectrum) | **0.47\*\*\*** | **-0.38\*\*** | **0.30\*** | 0.07 |
| Same pitch (lowpass) | - 0.07 | 0.07 | -0.11 | 0.11 |
| Different pitch (full spectrum) | **- 0.48\*\*\*** | **-0.32\*** | **-0.38\*\*** | 0.05 |
| Different pitch (lowpass) | -0.26 | 0.02 | -0.25 | 0.17 |

As shown by the scatterplot in the right panel of Fig. 8, the SRM magnitudes were distributed around zero indicating that lowpass filtering eliminated the benefit of spatial segregation between the target and the maskers. Consistent with our findings for the full-spectrum speech, Gallun et al. (2013) also reported a significant effect of age on SRM when using the same spatial locations for the target and maskers as the ones used here. A few studies found a significant effect of hearing sensitivity but not age on SRM in speech-on-speech masking despite applying amplification to spectral regions corresponding to hearing loss (Glyde et al., 2013; Glyde et al., 2015). Srinivasan et al. (2016) showed that for individuals with age-related hearing loss, age was a dominant predictor for SRM for smaller target/maskers spatial separations while the degree of hearing loss dominated SRM for larger spatial separations. This could explain the difference between the significant effects of age observed in this and the Gallun et al. (2013) study and significant effects of hearing loss but not age in the Glyde *et al.* (2013; 2015) studies, which used maskers located at -+90 deg. An apparent caveat with this explanation is that Srinivasan et al. (2016) found hearing loss to be a dominant predictor of SRM for target\maskers spatial separations above -+8 deg and age was a dominant predictor only for smaller spatial separations. However, listeners with hearing loss tested in that study had substantially greater loss of hearing sensitivity than the older listeners recruited in this study. In addition, reverberation was added to our speech stimuli, possibly shifting the spatial separations corresponding to the trade-off between the dominant roles of age and hearing loss toward larger spatial separations (Srinivasan et al., 2017).

As shown in Fig. 7, lowpass filtering of the stimuli reduced the amount of SRM for all age groups indicating significant contributions of frequency components above 2 kHz to spatial segregation of speech stimuli. This result is consistent with data from Kidd et al. (2010) showing that lowpass filtering reduces SRM and that significant SRM is observed for speech that is bandpass filtered into a high-frequency range between 3 and 6 kHz. It is possible that age effects on SRM were not seen for lowpass-filtered speech because of very small or absent SRM in many participants due to the limited speech spectrum.

## 5 Cognitive assessment

### 5.1 Stimuli and Procedure

All partcipants who completed the speech task were asked to complete two parts of a cognitive trail making test, [TMT; (Reitan, 1955)] implemented in the Psychology Experiment Building Language platform [PEBL version 2.0.4; (Mueller et al., 2014)]. In part A, participants were asked to connect 25 numbers in sequence, 1-2-3- and so on, and in part B, they were asked to connect alternating numbers and letters of alphabet, 1-A-2-B- and so on, as fast as possible. Both parts were completed on a desktop monitor using a computer mouse. Times needed to complete each test were recorded and used as scores. Scores from part A are known to be influenced by such cognitive functions as processing speed, psycho-motor control, and visual search, whereas scores from part B additionally index executive function (Bowie et al., 2006). The TMT test was selected because previous studies have shown that performance on this test accounted for significant amounts of variance in just detectable interaural time differences (Füllgrabe et al., 2015; Shehorn et al., 2020; Strelcyk et al., 2019) and variance in speech scores in a speech-on-speech task involving spatial target/masker segregation (Füllgrabe et al., 2015; Woods et al., 2013). Prior to test administration, participants completed shorter practice versions of each part.

### 5.2 Results and discussion

The times needed to complete the TMT-A test averaged within each of the three groups were, 24.25 s (SE = 1.54) for the young group, 27.19 s (SE = 1.54) s for the middle group, and 30.15 s (SE = 1.45) s for the older group. For the TMT-B test, the average times were, 28 s (SE = 1.7) for the young, 34.2 s (SE = 2.61) for the middle, and 37.32 s (SE = 2.34), for the older group. A repeated-measures ANOVA performed on scores from the two parts of TMT test, with TMT task (A and B) as the within-subjects factor and age group as the between-subjects factor, showed a significant effect of TMT task [F(1,54) = 50.58, p < 0.001] and a significant effect of age group [F(2,54) = 5.828, p = 0.005]. The interaction between the main factors was not significant [F(2,54) = 2.510, p = 0.091]. The post hoc analyses showed that the older group took significantly longer to complete the tests than the young group (p = 0.018), but there were no significant differences between the scores for the older and middle, and the middle and young groups (p > 0.05). As expected, participants took significantly longer to complete part B than part A of the test (p < 0.001). For all participants pooled across the three age groups, there were significant correlations between the TMT scores and age (part A: r = 0.332, p = 0.012; and part B: r = 0.315, p = 0.017) suggesting some decline in cognitive abilities of our participants with age.

## 6 Correlation and regression analyses

Data from 27 participants who completed all the experiments within this study were used to investigate the strength of associations between all the obtained measures using multiple correlations. Figure 9 shows a heat map representing values of the correlation coefficients. As indicated by the darker areas near the diagonal with white asterisks, measures from the same experiment but for different set of parameters were significantly correlated with one another while measures from different tasks were uncorrelated in most cases, even though, with the exception of the cognitive tests, the measures were expected to tap into the same general mechanisms that underlie temporal envelope processing. There were a few weak correlations between measures from different experiments. The scores from both parts of the cognitive TMT test were negatively correlated with EFR for one condition (a 0-dB modulation depth in quiet). All envITD thresholds were significantly correlated with SRM for the different-pitch low-pass-filtered speech condition but for full-spectrum speech, SRM was significantly correlated only with envITD thresholds for the 2-kHz AM tones presented in quiet.

**Fig. 9.**
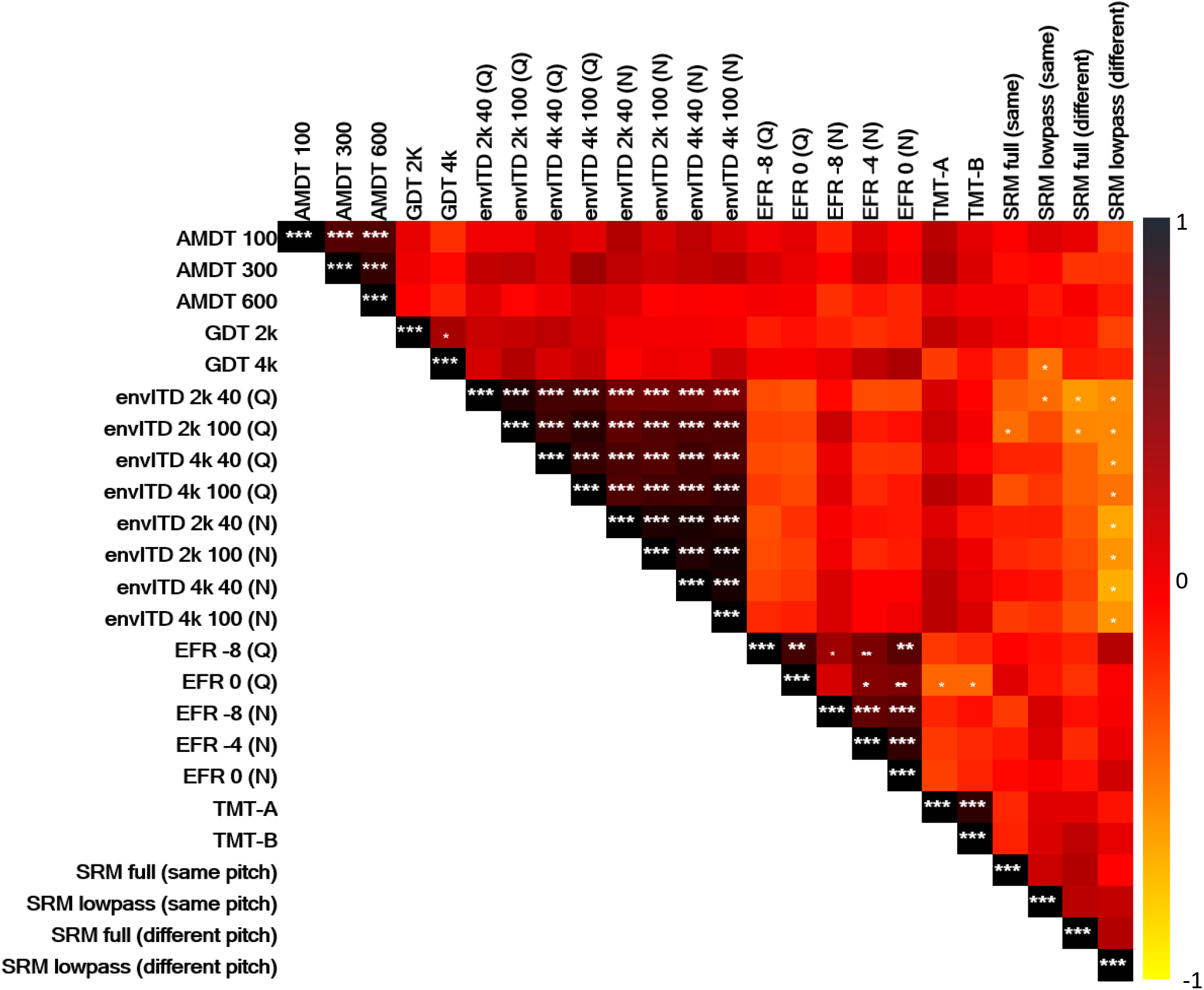
A heat map showing associations between different measures collected within this study. Asterisks denote significant correlations (*** p < 0.001, ** p < 0.01, and * p < 0.05). Asterisks in smaller font indicate significant correlations that do not survive corrections for multiple comparison.

A multivariate linear regression model was used to assess the proportion of variance in SRM explained separately and jointly by three variables, age, envITD for a 2-kHz carrier (average for the two modulation rates, 40 and 100 Hz), and ALLAVG. No other measures were entered into the regression model since they were not significantly correlated with SRM. Only the full-spectrum conditions were considered in the regression analysis because little or no SRM was observed with the lowpass-filtered speech for many participants. First, associations between each of the independent variables were tested using data from the 27 participants who completed all experiments within this study by using simple linear regressions. There was a significant correlation between participants’ age and ALLAVG (r = 0.48, p = 0.011), but the collinearity between the two variables was relatively weak so both variables were entered in the regression model. The average envITD thresholds for the 2-kHz carrier were not significantly correlated with age (r = 0.25, p = 0.21) or with ALLAVG (r = 0.22, p = 0.28).

For the same-pitch condition, all three independent variables were significant predictors of SRM, with age explaining 24.7% of the variance [F(1, 26) = 9.52, p = 0.005], envITD for a 2-kHz carrier explaining 16.1% of the variance [F(1, 26) = 5.97, p = 0.022], and ALLAVG explaining 14.5% of the variance [F(1, 26) = 5.43, p = 0.028] in the dependent variable. Jointly, the three variables explained 31.3% of the variance in SRM. With all three variables entered into the model, age was the dominant predictor (24.7% of the variance explained), with envITD and ALLAVG adding only 5.1% and 1.4% to the variance explained, respectively.

For the different-pitch condition, only two variables, age and average envITD threshold, were significant predictors of SRM. Age explained 42.3% of the variance [F(1,26) = 20.08, p < 0.001] and average envITD threshold explained 15% of the variance [F(1, 26) = 5.61, p = 0.026] in SRM when considered separately. Jointly, the two variables explained 47.9% of variance in SRM [F(2, 26) = 12.95, p < 0.001], with age being the dominant predictor (42.3%) and average envITD threshold adding only 5.6% of the variance explained. With the two variables entered, only age remained a significant predictor.

In summary, although the proportions of variance in SRM explained by the independent variables in both linear regression models were small, age emerged as the main predictor. Although the cognitive abilities evaluated in this study using two TMT tests declined with partcipants’ age, even for the sample of 27 individuals who completed all experiments in this study (part A: r = 0.40, p = 0.022, and part B: r = .42, p = 0.018), this decline could not explain the age-related reduction in SRM in any of the speech conditions since scores from TMT part A and B were not significantly correlated with SRM (Fig. 9).

## 7 General discussion

Aging is accompanied by changes to auditory processing that affect our ability to efficiently analyze complex auditory scenes, and crucially, the ability to process speech in noisy and reverberant environments. These changes include progressive loss of hearing sensitivity due to damage to hair cells and strial degeneration in the cochlea (Dubno et al., 2013; Ramadan et al., 1989; Schuknecht et al., 1993; Schulte et al., 1992; Wu et al., 2020), diffuse loss of synaptic connections between the cochlear hair cells and auditory-nerve fibers (Wu et al., 2020; Wu et al., 2019), abnormalities in neural coding in midbrain and cortical areas (Alain, 2014; Anderson et al., 2012; Presacco et al., 2016; Tremblay et al., 2003), and a decline in cognitive functions (Craik et al., 2011; Füllgrabe et al., 2015; Humes et al., 2013; Van der Linden et al., 1994; Ward et al., 2016). In this study, we focused on the effects of aging and hearing sensitivity on temporal envelope processing and on the role that changes in temporal processing may have on speech understanding. In addition to analyzing speech scores in a variety of conditions, we investigated age-related changes to speech recognition in “cocktail party” situation (Cherry, 1953) where better segregation of the target from background can be achieved by an effective use of spatial cues (Best et al., 2012; Gallun et al., 2013; Kidd Jr et al., 2016; Marrone et al., 2008; Swaminathan et al., 2015). SRM for small spatial segregation was used as a measure of the ability to use spatial cues.

It has been suggested that cochlear synaptopathy, shown to be a result of healthy aging (Sergeyenko et al., 2013) and excessive noise exposure (Kujawa et al., 2009) in animals, adversely affects processing of information carried by fast temporal changes in acoustic stimuli presented at medium to high levels in noise. In humans, direct evidence of cochlear synaptopathy has been provided by temporal bone studies (Makary et al., 2011; Viana et al., 2015; Wu et al., 2019). Despite extensive efforts, evidence consistent with the hypothesized adverse effects of cochlear synaptopathy on auditory perception in living humans remains elusive (Grose et al., 2017; Guest et al., 2017a; Johannesen et al., 2019; but cf. Liberman et al., 2016; Mepani et al., 2020; Prendergast et al., 2019; Prendergast et al., 2017a; Prendergast et al., 2018; Prendergast et al., 2017b; Shehorn et al., 2020; e.g., Spankovich et al., 2014). Early studies of cochlear synaptopathy tested mainly young normal-hearing listeners trying to avoid confounding effects of hearing loss that could explain the observed effects or offset them (Garrett et al., 2019). Because studies of human temporal bones provide direct evidence of increased cochlear synaptopathy with age (Wu et al., 2019), the approach taken in this study was to use participants with ages spanning a wide range and to separate the effects of age and hearing sensitivity on all the measures in statistical analyses. We found a significant correlation between age and average hearing sensitivity for all participants of this study (r = 0.48, p = 0.011). Although the correlation was not very strong, some portion of associations with age was inevitably removed by statistically controlling for differences in hearing thresholds.

### 7.1 Comparisons with other studies of age effects on envelope processing

Based on the hypothesized effects of cochlear synaptopathy, it was expected that performance in tasks that require robust processing of temporal information should decline with age after differences in hearing sensitivity have been accounted for and that results from different temporal-processing tasks should be significantly correlated with one another. The psychophysical and electrophysiological measures collected in this study do not support these expectations. Among the three behavioral tasks, only gap-detection thresholds were adversely affected by age. The age effect on the ability to detect gaps is consistent with previous findings that gap-detection thresholds increase with age (Helfer et al., 2009; Kumar et al., 2011; Schneider et al., 1999; Snell, 1997). AM-detection thresholds for fast modulation rates improved with age for two of the three modulation rates used in this study (100- and 300-Hz AM). The statistical analyses revealed that the improvement was associated with increased high-frequency hearing loss rather than with age per se. In contrast, data from most previous studies showed higher AM-detection thresholds in older than younger participants, for tonal and noise carriers, particularly for higher modulation rates (He et al., 2008; Kumar et al., 2011; Takahashi et al., 1992). Increased AM-detection thresholds were also found in listeners with hearing loss (Bacon et al., 1985). On the other hand, Prendergast et al. (2019) found significant improvement of AM-detection threshold for 25-Hz AM in a 4-kHz carrier with increasing hearing threshold at 4 kHz. It is worth noting that this and the Prendergast et al. (2019) study used much larger sample size than most earlier studies, and in this and Prendergat et al.’ s studies older participants had only mild-to-moderate hearing loss limited to high frequencies.

No significant effects of age or hearing sensitivity were found in the data from the binaural envITD task performed in quiet and in the presence of a masking noise, for two modulation rates, 40 and 100 Hz. Although for the faster AM rate, the noise significantly raised envITD thresholds compared to those in quiet, listeners across the three age groups, young, middle, and older, were equally affected by its presence. No effects of age and hearing sensitivity for adults with normal hearing and mild-to-moderate high-frequency hearing loss were also found by Prendergast et al. (2019). Inconsistent with expectations based on hypothesized effects of cochlear synaptopathy, no significant correlations between results from different behavioral tasks were observed (Fig. 9), even though performances for different parameters within each task were significantly correlated. This finding suggests that differences in specific tasks used to investigate the integrity of temporal processing can lead to different conclusions about effects of cochlear synaptopathy on this important auditory function. The outcomes of the behavioral experiments in this study are broadly consistent with findings from studies on cochlear synaptopathy that have used only young normal-hearing listeners (Grose et al., 2017; Prendergast et al., 2017a; Yeend et al., 2017) and a few other studies that also used older listeners with normal and near-normal hearing (Grose et al., 2019; Prendergast et al., 2019; Schoof et al., 2014).

The results of the electrophysiological EFR measurements also did not show a pattern consistent with that expected from cochlear synaptopathy. The ANOVA showed a significant effect of age overall at the two carrier frequencies tested, 2 and 4 kHz. However, post hoc pairwise comparisons showed that the PLVs were significantly reduced in older compared with younger adults only for fully (100%) modulated tones presented in quiet. Animal data suggest that cochlear synaptopathy mainly affects auditory nerve fibers with low and medium spontaneous rate (Furman et al., 2013). These fibers have been shown to reliably encode stimuli presented at medium to high levels in masking noise (Costalupes et al., 1984). Loss of peripheral synapses with age should therefore be most evident for the EFRs measured in noise. In quiet, high-spontaneous-rate fibers, less affected by synaptopathy, should reliably encode AM of tones in quiet, even in the presence of cochlear synaptopathy. Correlations between EFRs and age as well as HFAVG were not significant for any condition, consistent with the results reported by Prendergast et al. (2019). EFR PLV values were also not significantly correlated with any behavioral measures of temporal processing (see Fig. 9).

### 7.2 Speech-on-speech masking and predictors of spatial release from masking

Aging individuals often complain about difficulty understanding speech in noisy environments even when their hearing is normal by audiometric standards (Dubno et al., 2002; Helfer et al., 2008). The discovery of cochlear synaptopathy, a disorder that does not affect hearing sensitivity (Kujawa et al., 2009), provided a potential explanation for the reported age-related difficulties with speech processing. Understanding speech in the presence of other intelligible speech has been shown to be particularly challenging (Helfer et al., 2008; Rajan et al., 2008) and speech-on-speech masking is arguably most relevant to everyday listening challenges. Our results showed significant age effects in all conditions tested, i.e., both spatial locations of speakers and the same and different voice pitch, with full-spectrum speech (see Fig. 6). Average hearing sensitivity was not a significant factor in most conditions, possibly because our listeners had relatively mild hearing loss limited to high frequencies and because speech components were amplified in the regions of hearing loss.

A significant effect of age on speech recognition in two-talker babble was also found by Schoof et al. (2014) but Prendergast et al. (2019) reported no age effects. Differences in the design and particularly in speech material used for testing could contribute to the different outcomes. This study, like Schoof et al. (2014), used IEEE sentences which vary in syntactic content. Prendergast et al. (2019) used coordinated response measure, with sentences spoken by the target and competing speakers having the same syntactic structure. Using such a matrix-style speech material for both the target and maskers is problematic because it yields incorrect responses that result mainly from confusions rather than disruptions of speech understanding. Rennies et al. (2019) showed that over 60-80% errors for a matrix-style speech was due to participants reporting words that were spoken by the maskers and not the target.

SRM for full-spectrum speech, calculated from the difference between scores in non-colocated and colocated conditions, was significantly reduced in older listeners and the decline was already apparent in our middle-aged participants. The age affect was not due to differences in hearing sensitivity as shown by partial correlations in Table 4. The age-related decline in the ability to use spatial cues was uncorrelated with monaural measures of temporal processing but significant correlations were observed between SRM and envITD thresholds for a 2-kHz carrier. A multivariate regression analysis with SRM used as the dependent measure, and with independent variables of age, envITD threshold for a 2-kHz carrier averaged across two modulation rates used, 40 and 100 Hz, and average hearing sensitivity across the entire audiometric frequency range, showed that the main predictor of SRM was participants’ age. This was true for both full-spectrum conditions, with the target and maskers having the same voice pitch and with the maskers differing in voice pitch from the target. For the lowpass-filtered speech, age was not significantly correlated with SRM in either the same-pitch or different-pitch condition. However, SRM was not consistently observed across listeners with limited speech spectrum.

The significant age effects on speech recognition and on SRM could be due to cochlear synaptopathy although the lack of consistent age effects on different measures of temporal processing in this and other studies (Grose et al., 2019; Prendergast et al., 2019; Schoof et al., 2014) do not support this interpretation. It is possible that all measures are too variable across individuals and too strongly affected by a host of factors that are not related to temporal processing per se to reveal possibly small effects expected from synaptic loss in the auditory nerve (Oxenham, 2016). It is also possible that age-related deficits in temporal-fine-structure (TFS) processing (not measured here) are major contributors to difficulties with understanding the target in the presence of an interfering speech. Speech recognition scores measured using lowpass-filtered speech in this study were significantly affected by age even though all the listeners had clinically normal hearing in the region of all the speech components. A decline of TFS processing with age has been well documented (Füllgrabe, 2013; Grose et al., 2010; He et al., 2007) and the importance of TFS for speech recognition has been shown by a number of studies (Bernstein et al., 2011; Lorenzi et al., 2006; Moore, 2008; Pichora-Fuller et al., 2007; Schooneveldt et al., 1987). Schoof et al. (2014) found that detection of frequency modulation (FM) for a low-frequency carrier and low modulation rate was a significant predictor of speech recognition in two-talker babble, but not in steady-state or in amplitude-modulated noise. Detection of FM for the stimuli used by Schoof et al. (2014) is thought to reflect the fidelity of TFS processing (e.g., Moore et al., 1995; Moore et al., 1996; but cf. Whiteford et al., 2020). The adverse effect of age on the fidelity of TFS coding could be due to cochlear synaptopathy as synaptic loss in human temporal bones can be widespread over the entire length of the cochlea (Wu et al., 2019).

Speech processing is influenced by cognitive factors, such as selective attention (Best et al., 2007; Best et al., 2008; Clayton et al., 2016; Holmes et al., 2019), working memory (Akeroyd, 2008; Clayton et al., 2016), and processing speed (Füllgrabe et al., 2015; Woods et al., 2013). In this study, the scores from TMT tests that index processing speed showed significantly slower processing in older than younger participants. However, there was no correlation between SRM and performance on the TMT tasks. Likewise, Schoof et al. (2014) found no association between speech recognition in two-talker babble and processing speed or working memory, even though both were adversely affected by their participants’ age.

## 8 Final remarks

In summary, although significant age effects were observed for recognition of speech in two-talker babble with colocated and non-colocated speakers and for SRM, the psychophysical and electrophysiological measures provided no consistent evidence of deficits in temporal envelope processing or evidence of a relationship between temporal envelope processing and SRM. Cognitive measures of processing speed could not explain the significant age effects on speech recognition and SRM. Large individual variability in all measures could have contributed to the lack of significant associations between measures that are thought to reflect the same aspects of auditory processing. Even though age explained a significant proportion of variance in SRM in this study, it only explained ~33 – 42% of variance for full-spectrum speech. Finding all the major contributors to the variability in speech performance is crucial for understanding the deficits underlying poorer speech recognition in aging individuals. One way to overcome the large inter-subject variability is through longitudinal tracking relevant measures over different decades of life. Based on the results in this study, the speech-on-speech tasks with target and masker sentences that are low in context and do not follow the same syntactic structure combined with the use of speakers with similar voice pitches and small spatial separations appear to be good candidates for showing robust age effects. The challenge remains in finding robust measures that could best predict these age-related deficits. Based on our results, although temporal envelope plays an important role in speech understanding, temporal envelope processing does not appear to contribute to adverse effects of age on speech understanding in noisy and reverberant environments.

## ACKNOWLEDGMENTS

This work was supported by the NIH grant R01 DC015987 (M.W.). We would like to thank Alix Klang and Shashee Yang for assistance with data collection, and Anahita Mehta for input during the initial stages of setting up the EEG experiment.

## Notes

### Competing Interest Statement

The authors have declared no competing interest.

